# A whole-body atlas of non-graded BMP signaling activity in a sea anemone

**DOI:** 10.1101/2024.07.24.604959

**Authors:** Paul Knabl, David Mörsdorf, Grigory Genikhovich

**Affiliations:** Department of Neurosciences and Developmental Biology, University of Vienna, Austria; Vienna Doctoral School of Ecology and Evolution (VDSEE), University of Vienna, Austria

## Abstract

BMP signaling is responsible for the second body axis patterning in Bilateria and in the bilaterally symmetric members of the bilaterian sister clade Cnidaria – corals and sea anemones. However, medusozoan cnidarians (jellyfish, hydroids) are radially symmetric, and yet their genomes contain BMP signaling components. This evolutionary conservation suggests that BMP signaling must have other functions not related to axial patterning, which keeps BMP signaling components under selective pressure. To find out what these functions might be, we generated a detailed whole-body atlas of BMP activity in the sea anemone *Nematostella*. In the adult polyp, we discover an unexpected diversity of domains with BMP signaling activity, which is especially prominent in the head, as well as across the neuro-muscular and reproductive parts of the gastrodermis. In accordance, analysis of two medusozoan species, the true jellyfish *Aurelia* and the box jellyfish *Tripedalia,* revealed similarly broad and diverse BMP activity, supporting the versatile nature of the BMP pathway across anthozoan and medusozoan Cnidaria.

## Introduction

BMP signaling is initiated by the binding of dimeric BMP ligands to a tetrameric BMP receptor complex of two type II and two type I receptors. The complex recruits and phosphorylates the transcriptional effector SMAD1/5, which, as a result, binds the common-mediator SMAD4 and, as a trimeric complex, translocates to the nucleus to modulate BMP-responsive gene expression (**Figure 1A**). Since the initial discovery of BMPs in the context of bone and cartilage morphogenesis (Urist, 1965; Wu et al., 2024), BMP signaling has been studied in various vertebrate and invertebrate species, revealing far more versatile roles of the pathway. BMP signaling has been demonstrated to be involved in processes related to tissue homeostasis (Wang & Chen, 2018), apoptosis (Herrera et al., 2013; Kaltcheva et al., 2016; Zou & Niswander, 1996), cell fate commitment (Friedrichs et al., 2011; Hegarty et al., 2013), as well as gametogenesis, gonadogenesis and reproduction (Itman & Loveland, 2008; Lochab & Extavour, 2017; Regan et al., 2018). Moreover, BMP signaling is vital for instructing pattern formation of various tissues and organs, such as the limb bud (Vieira et al., 2019), the insect imaginal discs (Tripathi & Irvine, 2022), the gastrointestinal tract (Driver & Ohlstein, 2014; Zhang & Que, 2020), the nervous system (Hegarty et al., 2013) and, arguably most famously, the dorsoventral body axis across Bilateria (Bier & De Robertis, 2015; Morsdorf et al., 2024). While the roles of BMP signaling in Bilateria are multifaceted, its functions in non-bilaterian Metazoa are much less explored.

**Figure 1.**
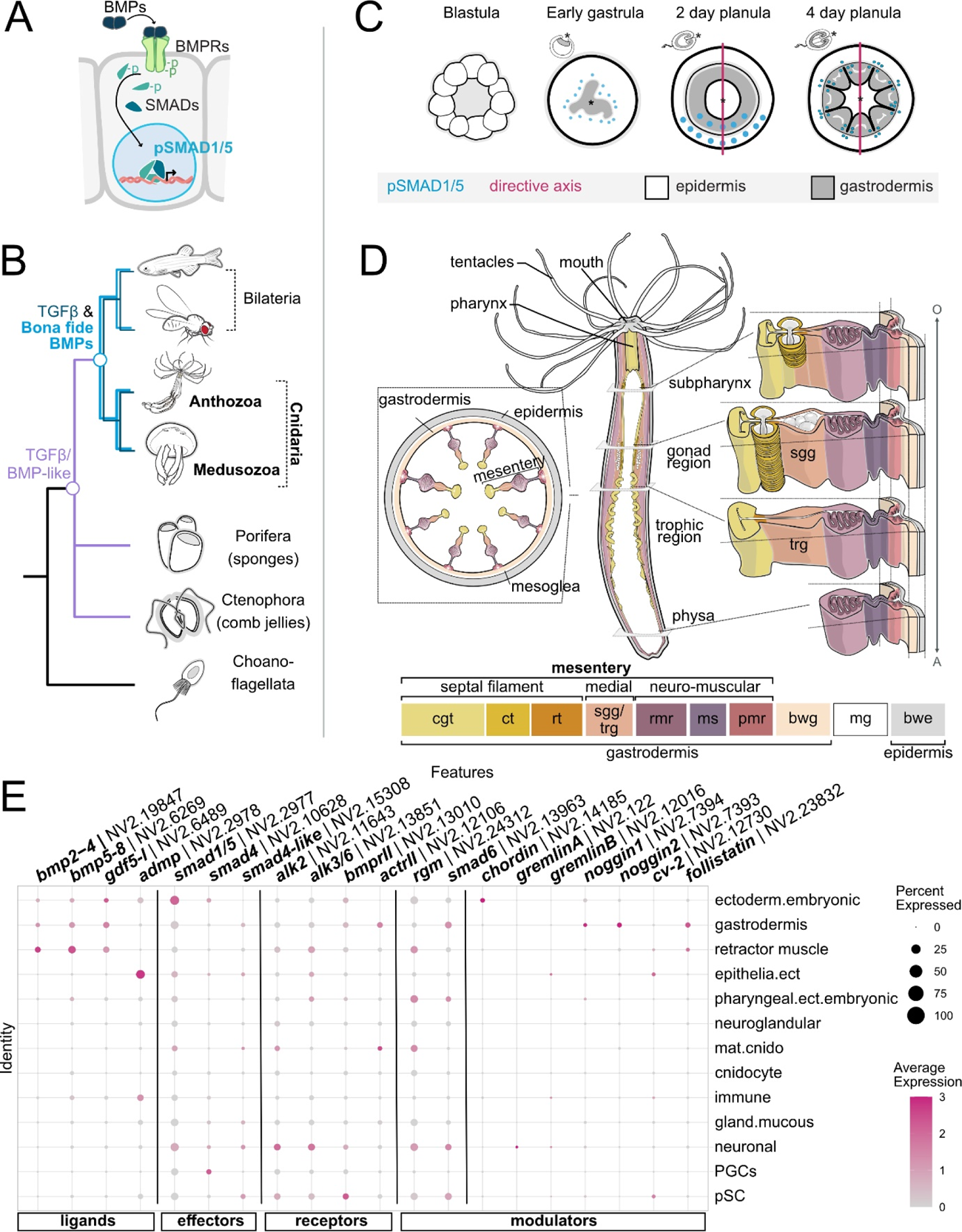
BMP pathway genes are evolutionary conserved between anthozoan Cnidaria and Medusozoa. (A) Activation of BMP signaling results in the phosphorylation of effector SMAD1/5 and its nuclear translocation to regulate BMP target gene transcription. (B) Evolutionary conservation of TGF-ß molecules is metazoan-specific, bona fide BMP pathway genes are conserved in Bilateria and Cnidaria (C) Dynamics of BMP signaling during early development of the sea anemone *Nematostella vectensis*. (D) Schematic of sea anemone *Nematostella* adult polyp highlighting mesentery anatomy at different body levels. (E) Seurat Dot plot visualizing the expression of BMP pathway genes in single-cell transcriptomic data of *Nematostella* adult polyp for coarse, tissue-level clustering (Cole et al., 2023; Cole et al., 2024; Steger et al., 2022). bwe - body wall epidermis, bwg - body wall gastrodermis, mg - mesoglea, pmr – parietal muscle region, ms – mesentery stalk, rmr - retractor muscle region, sgg – somatic gonad gastrodermis, trg – trophic gastrodermis, rt – reticulate tract, ct – ciliated tract, cng – cnidoglandular tract.

BMP pathway members are not exclusive to Bilateria but highly conserved in the bilaterian sister clade Cnidaria, a phylum comprising Anthozoa (sea anemones and corals) and Medusozoa (jellyfish and hydroids) (**Figure 1B**), the latter encompassing Hydrozoa (hydroids), Staurozoa (stalked jellyfish), Scyphozoa (true jellyfish) and Cubozoa (box jellyfish). All cnidarian classes possess the full complement of the intracellular BMP machinery, consisting of *bona fide* BMP ligands, receptors and effectors (Genikhovich & Technau, 2017; Gold et al., 2019; Kayal et al., 2018; Khalturin et al., 2019; Leclere et al., 2019; Lewis Ames et al., 2016; Ohdera et al., 2019). Most of our knowledge of BMP signaling to-date stems from Anthozoa, where it has been mainly studied in the sea anemone *Nematostella vectensis* and only in the context of axial patterning. Similar to Bilateria, *Nematostella* and other Anthozoa form a bilaterally symmetric body plan that is determined by a secondary, “directive” body axis and regulated by a BMP signaling gradient (Genikhovich et al., 2015; Knabl et al., 2024; Leclere & Rentzsch, 2014). In the embryo, graded BMP signaling was shown to be responsible for the formation and patterning of the directive axis, requiring the interaction of multiple BMP signaling components. Knockdown of BMP signaling pathway members including *bmp2/4, bmp5-8, gdf5-like* (=*gdf5-l*)*, chordin, gremlin* and *rgm* resulted in defects or complete loss of the directive axis and, consequently, impair patterning of the gastrodermal body layer^1^ (Genikhovich et al., 2015; Knabl et al., 2024; Leclere & Rentzsch, 2014; Saina et al., 2009). The dynamics of BMP signaling have been studied during early *Nematostella* development, using an antibody against the phosphorylated form of the transcriptional effector of BMP signaling, pSMAD1/5 (Genikhovich et al., 2015; Knabl et al., 2024; Leclere & Rentzsch, 2014). In the *Nematostella* late gastrula, pSMAD1/5 forms a bilaterally symmetric activity gradient, which disappears in the late planula larva (4 dpf = days post-fertilization) once the directive axis is established and patterned ((Knabl et al., 2024), **Figure 1C**). In the 4 dpf *Nematostella* planula, BMP signaling activity becomes radially symmetric and localizes to the oral epidermis, as well as in the newly formed gastrodermal folds termed the mesenteries ((Knabl et al., 2024), **Figure 1C**). To this point, it remains undetermined if and to which extent BMP signaling is required in other developmental processes beyond larval development of *Nematostella*.

Unlike bilaterally symmetric Anthozoa, all Medusozoa lack the directive body axis and feature a radially symmetric body plan. However, expression and transcriptomic data suggest that BMP signaling may be active in different medusozoan life stages, yet it is unknown which processes, if not the formation of a secondary axis, are regulated by BMP signaling in these animals. Previous analyses in several medusozoan species revealed generally broad expression profiles of BMP pathway genes across different body regions: The BMP ligand *bmp5-8* marks the tentacle zone in the polyp of the scyphozoan *Aurelia* and the hydrozoans *Hydra, Podocoryne* and *Clytia*, as well as the radial canals and sensory structures (rhopalia) in the *Aurelia* ephyra and in the medusae of *Clytia* and *Podocoryne* (Khalturin et al., 2019; Kraus et al., 2015; Reber-Muller et al., 2006; Reinhardt et al., 2004). In the hydroid *Hydractinia*, the expression of a putative *bmp receptor* was found in developing oocytes and the male gonophore gastrodermis (Sanders et al., 2014). To our knowledge, in Medusozoa, the activity of BMP signaling has not yet been directly addressed by visualizing phosphorylated SMAD1/5 in any published work.

Given that BMP signaling remains active in the sea anemone after directive axis formation and that BMP components are present in medusozoan species lacking the secondary axis, we aimed to determine what roles other than directive axis patterning may be regulated by BMP signaling in Cnidaria. In this work, we analyzed the activity of BMP signaling under homeostatic conditions in several cnidarian species, with a special focus on the adult polyp of *Nematostella*. We analyzed the publicly available scRNA-seq data (Cole et al., 2023; Cole et al., 2024; Steger et al., 2022) and found a broad expression of BMP signaling components across many cell types. Then, using an antibody against the activated form of the BMP effector SMAD1/5, we generated a morphological atlas of pSMAD1/5 activity in the whole adult polyp of *Nematostella*, highlighting the surprisingly diverse activity domains across different body regions, especially in the inter-tentacle epidermis and specific portions of the mesentery gastrodermis (**Figure 1D**). In the latter, we detected a population of pSMAD1/5-positive extra-epithelial cells that, however, was distinct from previously described vasa2-positive basiepithelial stem-like cells. Together with a first characterization of BMP signaling in the jellyfish *Aurelia* and *Tripedalia,* we find evidence for the activity of BMP signaling in multiple cell types across different body regions, suggesting versatile roles of BMP signaling in Cnidaria.

## Results

### Single cell RNA-seq data analysis suggests broad expression profiles of BMP pathway components across multiple cell populations in the adult *Nematostella* polyp

Thus far, expression studies of BMP pathway components and BMP signaling activity in *Nematostella* have been largely limited to early stages of development and the context of directive axis patterning (Finnerty et al., 2004; Genikhovich et al., 2015; Knabl et al., 2024; Leclere & Rentzsch, 2014; Matus, Pang, et al., 2006; Matus, Thomsen, et al., 2006; Rentzsch et al., 2006; Saina et al., 2009). To get some information about where we can expect BMP signaling in adult polyp, we interrogated the publicly available single-cell transcriptomic datasets for the expression of the BMP pathway genes (Cole et al., 2023; Cole et al., 2024; Steger et al., 2022). Here, we focused on datasets of female and male polyps (“adult tissues”) with a coarse clustering approach recovering 13 distinct clusters based on the study by Cole et al. 2024 (**Figure 1E**). The *Nematostella* genome contains gene orthologues for four BMP ligands (*bmp2/4*, *bmp5-8*, *gdf5-l* and *admp*), four BMP receptors (two type I receptors, *alk2* and *alk3/6*, and two type II receptors, *bmprII* and *actRII*) and two BMP effectors (*smad1/5* and *smad4*, as well as a “truncated” *smad4-like*). In addition, several orthologues of potential BMP signaling antagonists are present, including transmembrane (rgm), intracellular (*smad6*) and extracellular or secreted molecules (*chordin*, *gremlinA*, *gremlinB*, *noggin1, noggin2*, *crossveinless-2* and *follistatin*).

BMP receptors and BMP effectors displayed low expression levels but with surprisingly broad expression profiles and transcripts detectable in multiple cell clusters. Intracellular components were present in mature cnidocytes (“mat.cnido”), “retractor muscle”, “gastrodermis”, epithelial ectoderm (“epithelia.ect”) and embryonic ectoderm (“ectoderm.embryonic”), while they were upregulated in putative stem cell (“pSC”), and neurosensory and -secretory (“neuronal”) cell types. In contrast, the expression of BMP ligands was spatially segregated, with *bmp2/4*, *bmp5-8* and *gdf5-l* exhibiting similar expression profiles in the “gastrodermis”, “retractor muscle” and “ectoderm.embryonic”, whereas *admp* expression was restricted to the “epidermis”. For the expression of BMP receptor types, no obvious cell type-specific segregation was apparent. Together, the expression of BMP pathway genes in the single-cell transcriptomic data suggest that BMP signaling is potentially active in various cell populations, involving few “BMP secreting” cell types that can signal to a wider range of “BMP receiving” cell types. BMP pathway genes display low expression levels and are limited to a low number of cells per cluster (transcripts expressed in < 16% of cells of each cluster), which might be due to diverging expression profiles within the different clusters. Indeed, expression across the 91 adult-restricted transcriptomic states shows that the expression of BMP pathway genes can vary between different sub-clusters in each coarse cluster (**Fig. 1 - Suppl. 1**).

### BMP signaling forms distinct activity domains in the inner and outer epithelial layers of the adult polyp

To explore BMP signaling unrelated to axial patterning, we analyzed the activity of BMP signaling in the sexually mature polyp (**Figure 2A**), utilizing the cross-reactive commercial antibody against the phosphorylated form of the BMP signaling effector SMAD1/5, which we have successfully used in *Nematostella* embryos previously (Knabl et al., 2024). In the adult, we detected distinct domains of pSMAD1/5 activity in both epithelial body layers (**Figure 1D**), comprising the “outer” epidermis and the “inner” gastrodermis, which are separated by an extracellular matrix layer (mesoglea). Most of the epidermal epithelium was free of pSMAD1/5 activity, except for the oral domain (**Figure 2B**). Here, BMP signaling was detectable at the inter-tentacular spaces (**Figure 2C-E’, Figure 2 – Figure Suppl. 1**), as well as in longitudinal, epidermal stripes (**Figure 2I-L’, Figure 2 – Figure Suppl. 1)**. BMP signaling between the tentacles was present at each tentacle base with pSMAD1/5 activity mostly evident in the epidermis (**Figure 2C-E’**), slightly reduced in the gastrodermis (**Figure 2D**), and completely absent from other parts of the tentacles, except for the unspecific, non-nuclear staining of the capsules and threads of the tentacle-specific cnidocyte type – the spirocytes **(Figure 2 – Figure Suppl. 2)**. In the heterozygous *bmp2/4::mCherry* transgenic animals, in which mCherry reporter is expressed under control of the 4.5 kb regulatory region upstream of the *bmp2/4* start codon (Mörsdorf et al., 2024; Schwaiger et al., 2014), strong mCherry expression marks one side of the pharynx forming the ciliated groove, the siphonoglyph (**Figure 2F-H, Figure 2 – Figure Suppl. 3**). Previous work also showed that *bmp2/4* mRNA is present in the siphonoglyph cells of the planula larva (Finnerty et al., 2004), however we did not detect above-background pSMAD1/5 staining in this area in the adult.

**Figure 2.**
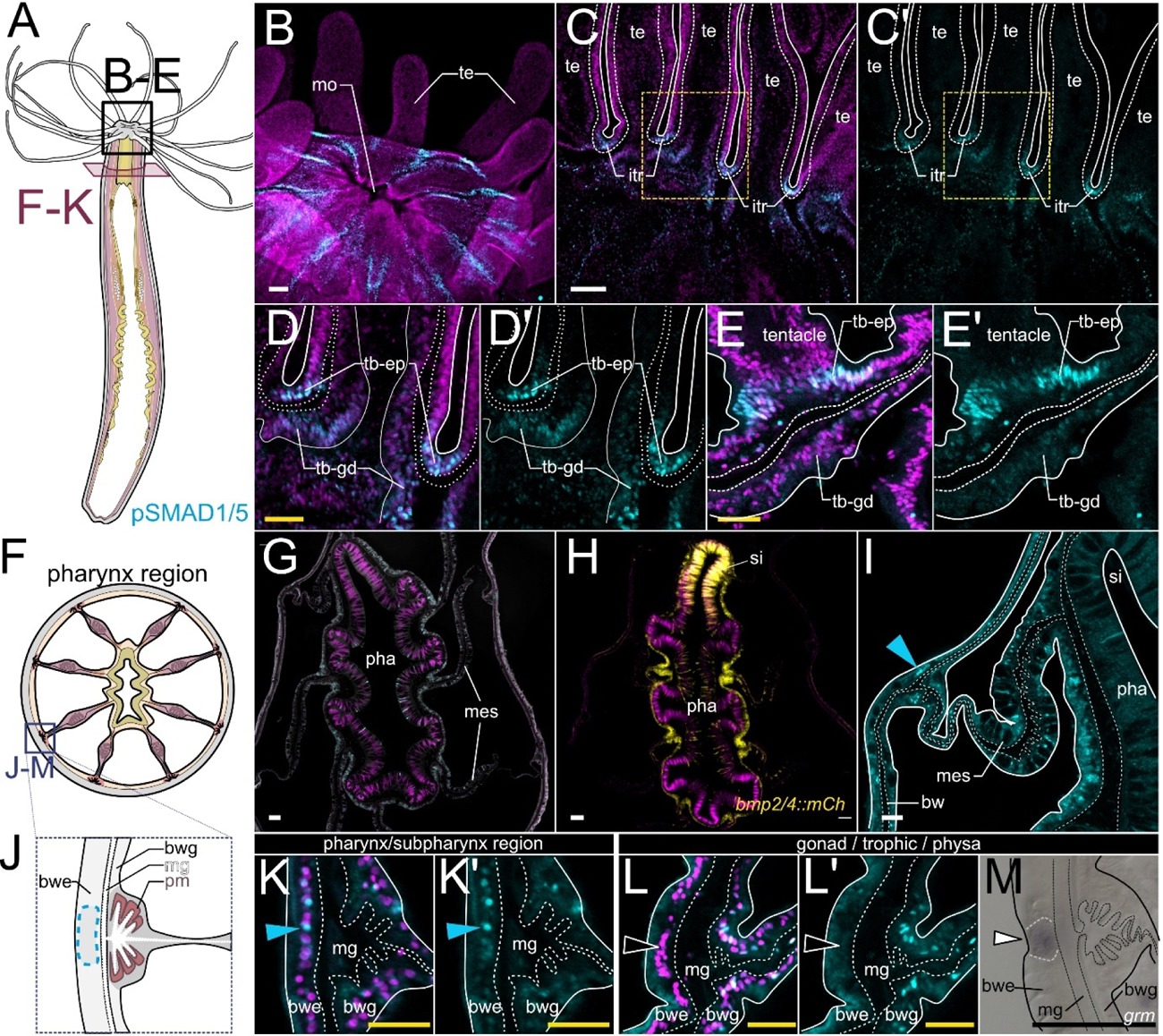
Epidermal BMP signaling is confined to the oral epithelium. Characterization of BMP signaling in the oral region by pSMAD1/5 (cyan) and DNA (magenta), outlines of tissue layers (solid white lines) and mesoglea (dashed white lines) are indicated. (A) mouth region (black box), cross-section of the body column at the pharynx level (red box), (B) pSMAD1/5 staining in the region of the hypostome (C) longitudinal cross-section of the tentacle and tentacle base (D-E) details of the tentacle base (F) schematic of a cross-section at the pharynx level (G) overview of a pharynx cross-section (H) *mCherry* expression in the bmp2/4::mCh reporter line highlights the ciliated groove (siphonoglyph) on one side of the pharynx (I) detail of a pharynx cross-section, pSMAD1/5-positive cells are located in the epidermis at the position of the mesenteric attachment site (blue arrow), (J) schematic of a mesenteric attachment site (K-K’) pSMAD1/5-positive nuclei in the epidermis of the mesenteric attachment site at the level of the pharynx and subpharynx (L-L’) BMP signaling is absent in the respective regions at the level of the gonad, trophic or physa region (M) epidermal *gremlin* expression. mo - mouth, te - tentacle, itr - inter-tentacular region, tb - tentacle base, tb-ep tentacle base epidermis, tb-gd - tentacle base gastrodermis, pha - pharynx, si – siphonoglyph, mes - mesentery, mg - mesoglea, bwe - body wall epidermis, bwg - body wall gastrodermis; scale bars 50 µm (white), 25 µm (yellow) and 100 µm (purple)

In the body wall epidermis, at the level of the pharynx, epidermal clusters of pSMAD1/5-positive cells assemble into eight longitudinal stripes (**Figure 2I-L’**), correlating with the positions of each mesentery attachment site in the gastrodermis (**Figure 2J**). Notably, discernible BMP signaling activity in the stripes was restricted to the pharyngeal and subpharyngeal region (**Figure 2K-K’**), while absent from more aboral positions (**Figure 2L-L’**). There, at the levels of the gonadal, trophic and physa region, BMP antagonist *gremlin* was expressed in the epidermis *vis-à-vis* the mesenterial attachment sites, and this area was free of pSMAD1/5 activity (**Figure 2M**).

In contrast to the orally restricted BMP signaling in the epidermis, BMP signaling in the gastrodermis was evident along the entire primary, oral-aboral (OA) body axis (**Figure 3**), and especially prominent in the mesenteries - the eight folds of the gastrodermal epithelium partitioning the gastric cavity (**Figure 1D**, **Figure 3A**), where pSMAD1/5 activity was highly pronounced. We did not observe any differences in pSMAD1/5 staining between different mesenteries, however, when comparing BMP signaling in the basal, medial and apical portion of the mesentery, we observed differences in activity depending on the position along the OA body axis (**Figure 3A-D’**). BMP signaling in the basal part of the mesentery (neuro-muscular mesentery, closest to the body wall) was the most consistent along the OA axis, with continuous and pronounced signaling activity from pharynx to physa (**Figure 3A-C’**, blue solid arrowheads). The apical domain of the mesentery (septal filament, cnidoglandular tract) displayed little nuclear pSMAD1/5, but often showed unspecific cytoplasmic or vesicular signal. Non-specific signal was repeatedly observed in mucoglandular cells that are most abundant in the trophic region (**Figure 3C-C’**, white arrowheads in the sf region**; Figure 2 – Figure Suppl. 2**). Unlike in the basal and apical domain, there were distinct differences of pSMAD1/5 domains in the medial mesentery portion, changing according to the body region along the primary body axis (**Figure 3A-D’,** mm region). While BMP signaling was broadly active in the medial mesentery of the pharynx and upper subpharynx region (**Figure 3B-B’**, solid blue arrowheads in the mm region), pSMAD1/5 was reduced or absent from the medial gastrodermis at lower positions (lower subpharynx region, gonad region and trophic region) (**Figure 3C-D’**, hollow white arrowheads in the mm region).

**Figure 3.**
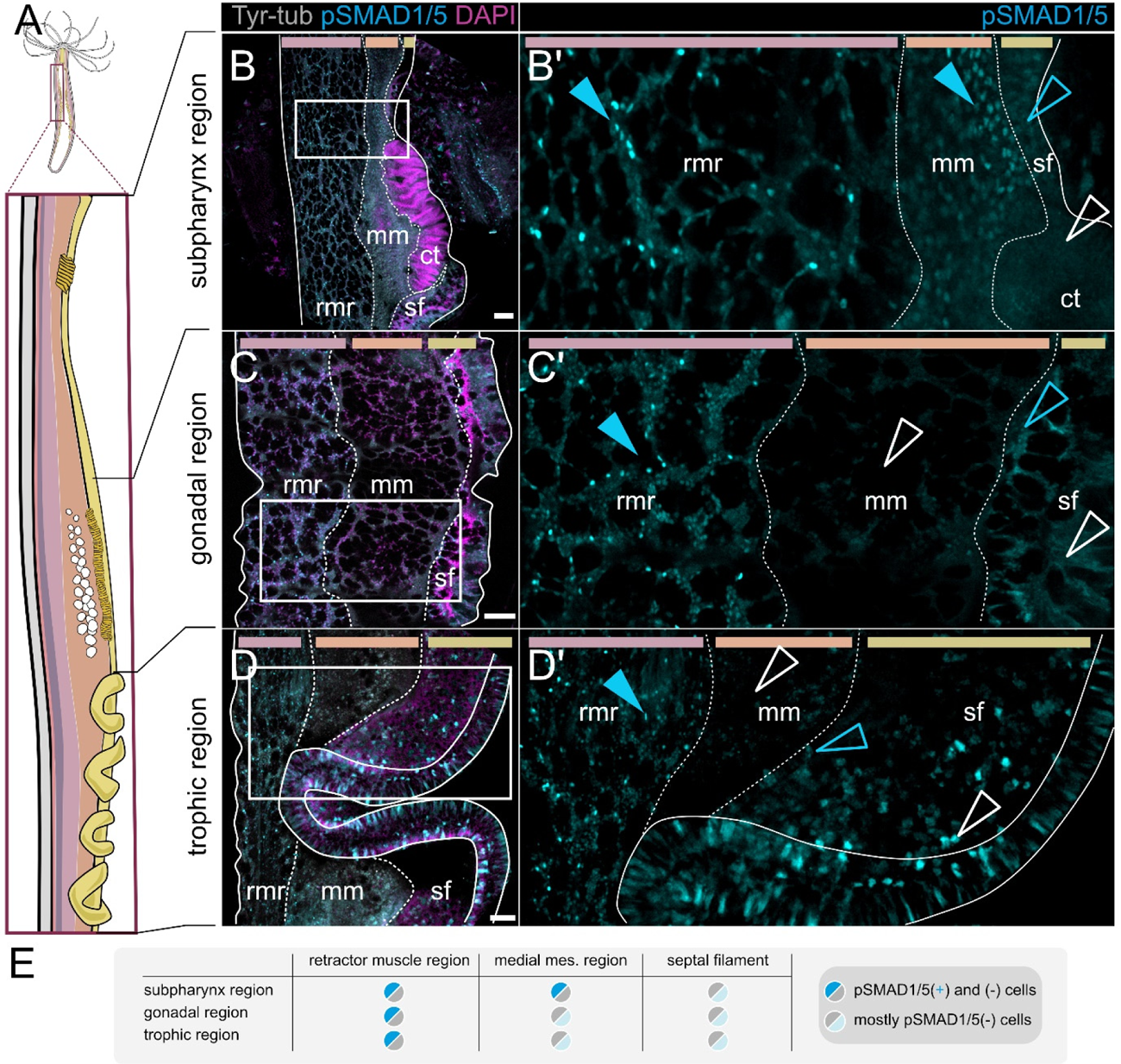
Different regions of the oral-aboral axis display different patterns of BMP signaling activity in the mesentery. (A) Sketch of the lateral overviews of mesenteries at different positions along the oral-aboral body axis shown on (B-D’). White lines demarcate different mesentery domains, pSMAD1/5 (cyan) and DNA (magenta). (B) subpharyngeal region displays pSMAD1/5 activity in the rmr (retractor muscle region) and mm (medial mesentery) (blue solid arrowheads), and few pSMAD1/5-positive nuclei in the cnidogladular tract of the septal filament (sf; blue, hollow arrowhead) (C) gonadal region and (D) trophic region, display pSMAD1/5 activity in the rmr, pSMAD1/5 activity is mostly absent in the mm (white, hollow arrow) and few pSMAD1/5-positive nuclei in the sf (blue, hollow arrowheads). (E) Summary of BMP signaling domains across different mesentery regions. Scale bars 50 µm. rmr - retarctor muscle region, mm – medial mesentery, sf – septal filament, ct – ciliated tract.

### Gastrodermal BMP signaling occurs in distinct domains along the basal-apical axis of the mesentery

To get a more detailed picture of BMP signaling in different parts of the mesentery, we analyzed the pSMAD1/5 activity in the individual mesentery domains, starting at the body wall and the basal part of the mesentery (neuro-muscular domain, **Figure 4**), moving along the medial mesentery domain and into the septal filament at the mesentery tip pointing towards the center of the gastric cavity (**Figure 5**). In the mesenteries, we could differentiate between (1) pSMAD1/5-positive epithelial cell populations in different domains of the mesentery and (2) pSMAD1/5-positive interstitial or basiepithelial cells, located in the extracellular matrix (mesoglea).

**Figure 4.**
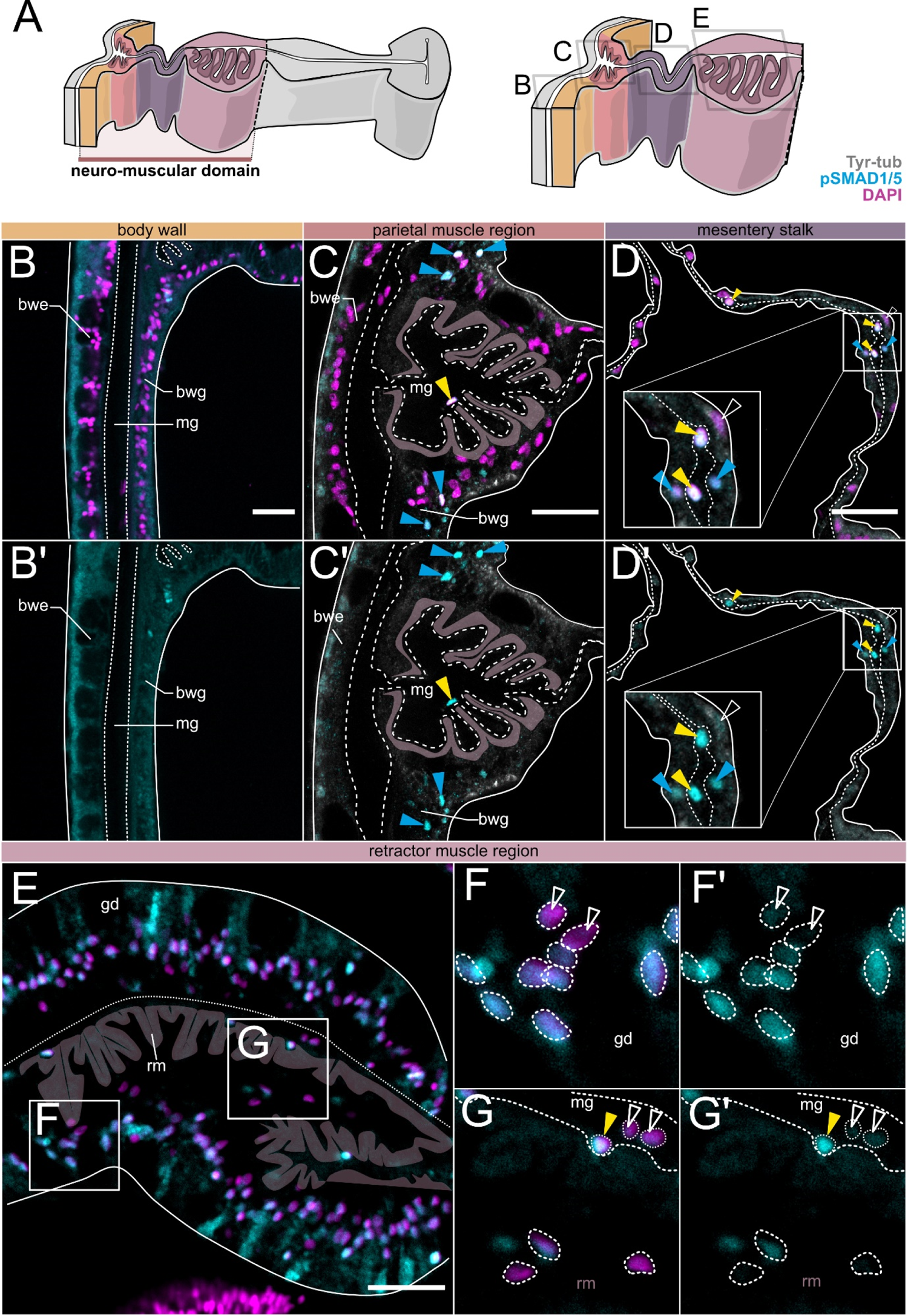
BMP signaling is active in epithelial and basiepithelial cells of the neuro-muscular mesentery. Immunostaining for pSMAD1/5 (cyan), Tyr-tubulin (grey) and DNA (magenta). (A) Schematic overview of a mesentery cross-section illustrating the morphological domains, highlighted in color are the basal mesentery regions. Images on (B-E) are transverse sections corresponding to regions boxed on (A). (F-G’) are high magnification views of areas boxed on (E). (B-B’) pSMAD1/5-negative body wall gastrodermis and epidermis. (C-C’) pSMAD1/5-positive epithelial cell clusters (blue arrowheads) and individual basiepithelial cells (yellow arrowheads.) (D-D’) pSMAD1/5-positive and -negative cells in the mesentery stalk in the epithelium and the mesoglea (yellow arrowheads). (E) cross-section of the retractor muscle region with pSMAD1/5-positive and pSMAD1/5-negative epithelial and basiepithelial cells, retractor muscle myonemes are highlighted in mauve. (F’-F’’) Magnified view of the retractor muscle region epithelium with pSMAD1/5-positive and-negative gastrodermal cells. (G’-G’’) visual section of the retractor muscle region with pSMAD1/5-positive and -negative epithelial and basiepithelial cells. Scale bars 25 µm. mg - mesoglea, bwe - body wall epidermis, bwg - body wall gastrodermis, gd – gastrodermis, rm – retractor muscle.

**Figure 5.**
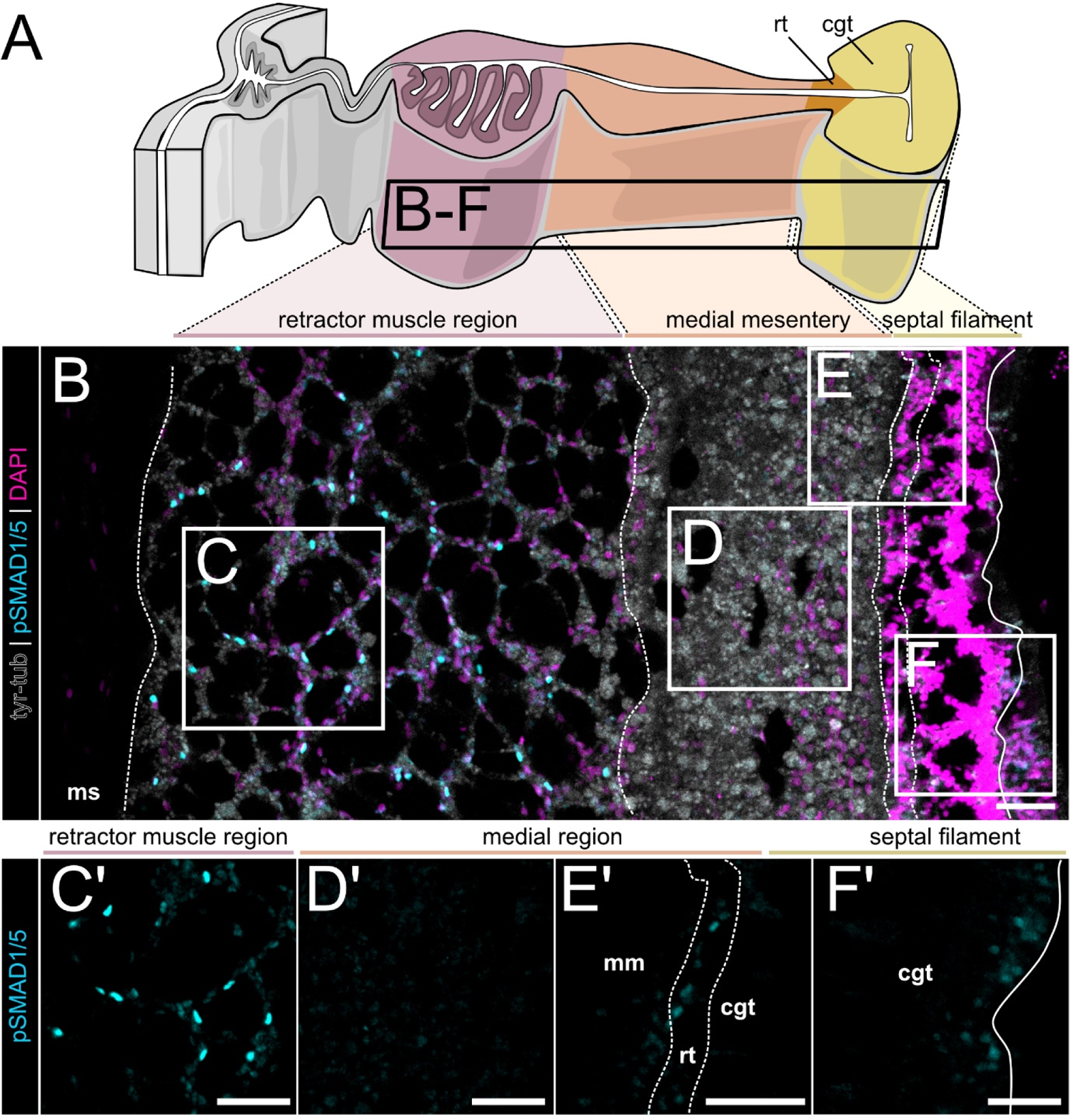
Differences in the BMP signaling domains of the medial and distal mesentery region. Immunostaining for pSMAD1/5 (cyan), Tyr-tubulin (grey) and DNA (magenta). (A) Schematic overview of a mesentery cross-section illustrating morphological domains, highlighted are the retractor muscle region and medial and distal mesentery domains. (B) lateral view of the mesentery at the level between subpharynx and gonad region. (C-C’) broad pSMAD1/5 activity in the retractor muscle region. (D-D’) pSMAD1/5 is mostly absent from the medial mesentery. (E-E’) individual pSMAD1/5-positive cells at the boundary of the medial mesentery and the septal filament in the reticulate tract. (F-F’) individual pSMAD1/5-positive cells in the cnidoglandular tract of the septal filament. rmr - retractor muscle region, mm - medial mesentery, rt – reticulate tract, cgt - cnidoglandular tract. Scale bars 50 µm.

The body wall gastrodermis between the mesenteries was generally free of BMP signaling (**Figure 4B-B’**), although individual pSMAD1/5-positive cells were occasionally observed. At the base of each mesentery, BMP signaling was active in two epithelial cell clusters located on both sides of the parietal muscle (**Figure 4C-C’**, blue arrowheads), while other cells of the parietal muscle region were pSMAD1/5-negative. The adjoining mesentery stalk contained pSMAD1/5-positive and -negative epithelial cells, distributed throughout the stalk (**Figure 4D-D’**). The gastrodermal epithelium of the retractor muscle region displayed pronounced pSMAD1/5 activity, also with intermixed communities of pSMAD1/5-positive and pSMAD1/5-negative cells (**Figure 4E-G’**).

Aside from epithelial cell populations, we also observed interstitial, basiepithelial cells that were present in the mesoglea of the mesentery and the body wall, across different locations of the body column, as previously described by others (Miramón-Puértolas & Steinmetz, 2023; Tucker et al., 2011). Notably, many basiepithelial cells located in the parietal muscle region, along the mesentery stalk and in the retractor muscle region were pSMAD1/5-positive (**Figure 4C-D’, G-G’**, yellow arrowheads), although we observed basiepithelial cells without BMP signaling activity in the same region as well. Strikingly, BMP signaling activity in these cells appeared to be restricted to the neuromuscular region of the mesentery but not to the other portions of the mesentery or other locations of the body column. This indicates that there are different subpopulations of basiepithelial cells present in the mesoglea that can be either pSMAD1/5-positive or -negative, and/or that BMP signaling in these cells is only active in a specific context, which may possibly be linked to their differentiation choices.

In the medial and apical mesentery (**Figures 5A, Figure 5 – Figure Suppl. 1**), activity domains of BMP signaling were less pronounced compared to the basal mesentery (**Figure 5B-F’**). In the medial domain, BMP signaling was mostly absent (**Figure 5B, D’-E’**), except at the levels of the pharynx and subpharynx (**Figure 3A-A’**), as described above, and in a narrow stripe of cells along periphery of the medial domain at its border to the reticulate tract (rt) in the gonadal region or to the cnidoglandular tract (cgt) in the trophic region (**Figure 5E’**). Other exceptions, observed in the somatic gonad, are outlined in a separate section below. The cnidoglandular tract was mostly devoid of BMP signaling, however, sporadically, pSMAD1/5-positive cells were also observed in different locations of the septal filament (**Figure 5F’, Figure 5 - Figure Supplement 1**).

Overall, we observed intermixed communities of pSMAD1/5-positive and -negative cells, with epithelial or basiepithelial localization, that form multiple BMP signaling domains in the mesentery. pSMAD1/5 activity domains of the basal, neuro-muscular mesentery are highly pronounced, compared to the more reduced or completely absent activity in the medial and apical mesentery. This heterogeneity in composition and location of pSMAD1/5 domains suggest that BMP signaling activity is not limited to a single BMP-responsive cell population but rather can be active in multiple cell types or cell states, which is supported by the broad expression of intracellular BMP components in the single-cell transcriptomic data.

### Medial and apical mesentery domains display reduced pSMAD1/5 activity but increased BMP expression

Our characterization of mesenteric BMP signaling domains revealed the intriguing reduction of signaling activity in the medial and apical mesentery including the gonad region (**Figure 6A-B**) that we aimed to address in more detail (**Figure 6, Figure 6 – Figure Supplement 1**). First, we analyzed the expression of BMP receptors using colorimetric *in situ* hybridization, and detected *alk2*, *alk3/6* and *actrII* but no *bmprII* mRNA in parts of the septal filament, with an upregulated expression in the reticulate and ciliated tracts in the gonad region of the polyp (**Figure 6C-F**). We also detected BMP signaling activity there, although the pSMAD1/5 staining appeared weaker than in the areas described in the previous chapters. In the septal filament, BMP signaling was active in the epithelial cells of the reticulate tract (**Figure 6G-G’**, blue solid arrowheads), while it was absent from basiepithelial cells. Similarly, low levels of BMP signaling were sometimes detectable in the ciliated tract of the gonad region (**Figure 6H-I’**), but often completely absent (**Figure 6G-G’**, white hollow arrowheads), and BMP signaling activity was never observed in the ciliated tract of the subpharynx region. Active BMP signaling in the ciliated tract seemed to be localized to the cells at the top rather than at the bottom of the folds (**Figure 6I-I’**). It remains unclear if these differences in the detection and localization of pSMAD1/5 signals reflect dynamic changes of BMP signaling in the ciliated tract or they are due to technical limitations of the antibody staining.

**Figure 6.**
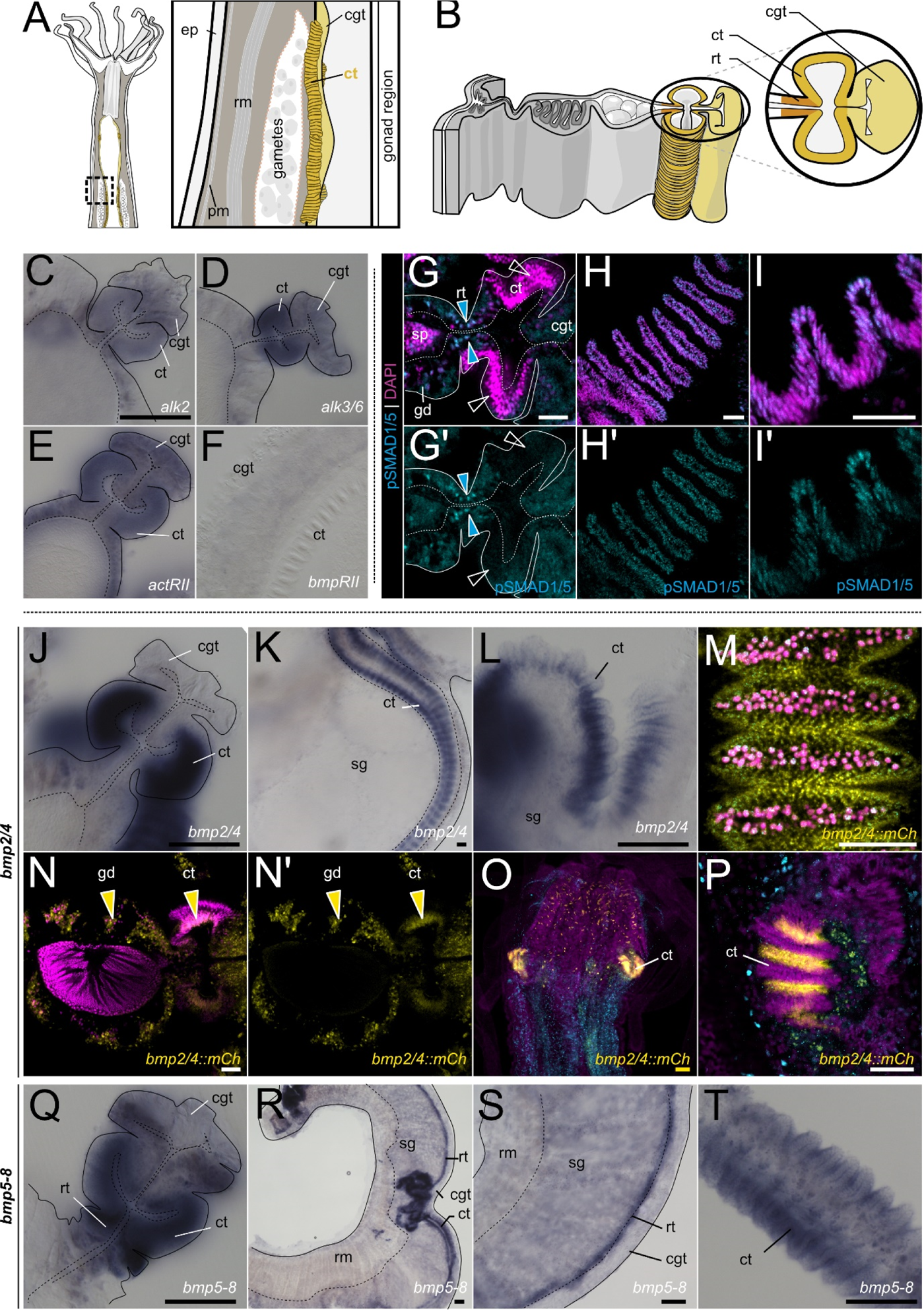
pSMAD1/5 activity, BMP and BMP receptor expression in the septal filament. (A-B) Schematics of the mesentery morphology at the level of the gonad region in a lateral view (A) and a transverse section (B). Sub-regions of the septal filament highlighted in shades of yellow. (C-F) BMP receptor expression of the type I receptors *alk2* (C) and *alk3/6* (D), and of the type II receptors *actRII* (E) and *bmprII* (F) in the septal filament. (G-I’) immunostaining for pSMAD1/5 (cyan) and DNA (magenta) in the septal filament. (G-G’) cross-section of the mesentery of a male in the gonad region, ciliated tracts are indicated (ct), blue arrowheads highlight pSMAD1/5 in the reticulate tract (rt), (H-H’) top view of the ciliated tract showing pSMAD1/5 activity in its folds. (I-I’) longitudinal optical section of the ciliated tract showing pSMAD1/5 activity in the folds. (J-L) *bmp2/4* expression in the septal filament in cross-section and lateral views. (M-O) mCherry expression in the *bmp2/4::mCherry* reporter line. (M-N’) mCherry is expressed in the mesentery gastrodermis and in the ciliated tract of the adult polyp. (O-P) *bmp2/4::mCherry* activity in the ciliated tract in the juvenile polyp. (Q-T) *bmp5-8* expression in the mesentery gastrodermis, reticulate tract and ciliated tract, cross-section and lateral views. Scale bars 25 µm (white), 50 µm (yellow) and 100 µm (black). ep – epidermis, pm – parietal muscle, rm - retractor muscle, gd – gastrodermis, sp – spermary, rt - reticulate tract, ct - ciliated tract, cgt - cnidoglandular tract, sg – somatic gonad.

In line with the expression of the BMP receptors and pSMAD1/5 immunoreactivity (**Figure 6C-F; H-I’**), in situ hybridization analysis also showed the expression of BMP ligand genes in the septal filament. We found *bmp2/4* (**Figure 6J-P**) and *bmp5-8* (**Figure 6Q-T**) expression in the ciliated tract, curiously, at the bottom of the folds, *i.e.* in a domain complementary to the areas of pSMAD1/5 activity at the top of the ciliated track folds. Unlike *bmp2/4*, in addition to the ciliated tract expression, *bmp5-8* is also strongly expressed in the reticulate tract of the septal filament (**Figure 6Q-S**). In agreement with in situ hybridization data, transgenic adult polyps of the *bmp2/4::mCh* reporter line displayed *mCherry* expression in the mesenterial gastrodermis and at the basis of the ciliated tracts, both in the subpharynx and the gonadal region (**Figure 6M-P, Figure 2 – Figure Supplement 3**).

In the gonads (**Figure 7A-B**), transcripts of BMP ligands *bmp2/4, bmp5-8* and *gdf5-like*, and BMP receptors *alk2*, *alk3/6*, *actrII* and *bmprII* were detectable in the gastrodermis of the somatic gonad (**Figure 7C-I’**), and several of the BMP signaling components were deposited in maturing gametes. In female polyps, in situ hybridization suggested maternal deposition of mRNAs of *bmp5-8*, *alk2*, *alk3/6*, *actrII*, and *bmprII* but not of *bmp2/4* or *gdf5-like* (**Figure 7C-I**). This was further supported by the RNA-Seq data deposited in the NvERTx database (Fischer & Smith, 2012; Helm et al., 2013; Warner et al., 2018), detecting maternal transcripts of *bmp5-8*, *alk2*, *alk3/6*, *actrII* and *bmprII* in unfertilized *Nematostella* eggs. According to the NvERTx, low *bmp2/4* and *gdf5-like* appear to be also provided maternally, however at much lower levels than *bmp5-8* and undetectable by in situ hybridization (**Figure 7 – Figure Supplement 1**). Curiously, maternal deposition of BMP components seems redundant, since active BMP signaling only becomes detectable at gastrulation (∼20hpf; (Knabl et al., 2024)). In situ hybridization analysis of the male polyp showed expression of *bmp2/4*, *bmp5-8*, *gdf5-l*, *alk2*, *alk3/6* and *actrII* in the somatic gonad, but no detectable staining of BMP RNAs in the germ cells. In contrast, mRNA of the BMP receptor *alk2* appears to accumulate in the periphery of the spermaries (i.e. in spermatogonia) (**Figure 7C’-I’**).

**Figure 7.**
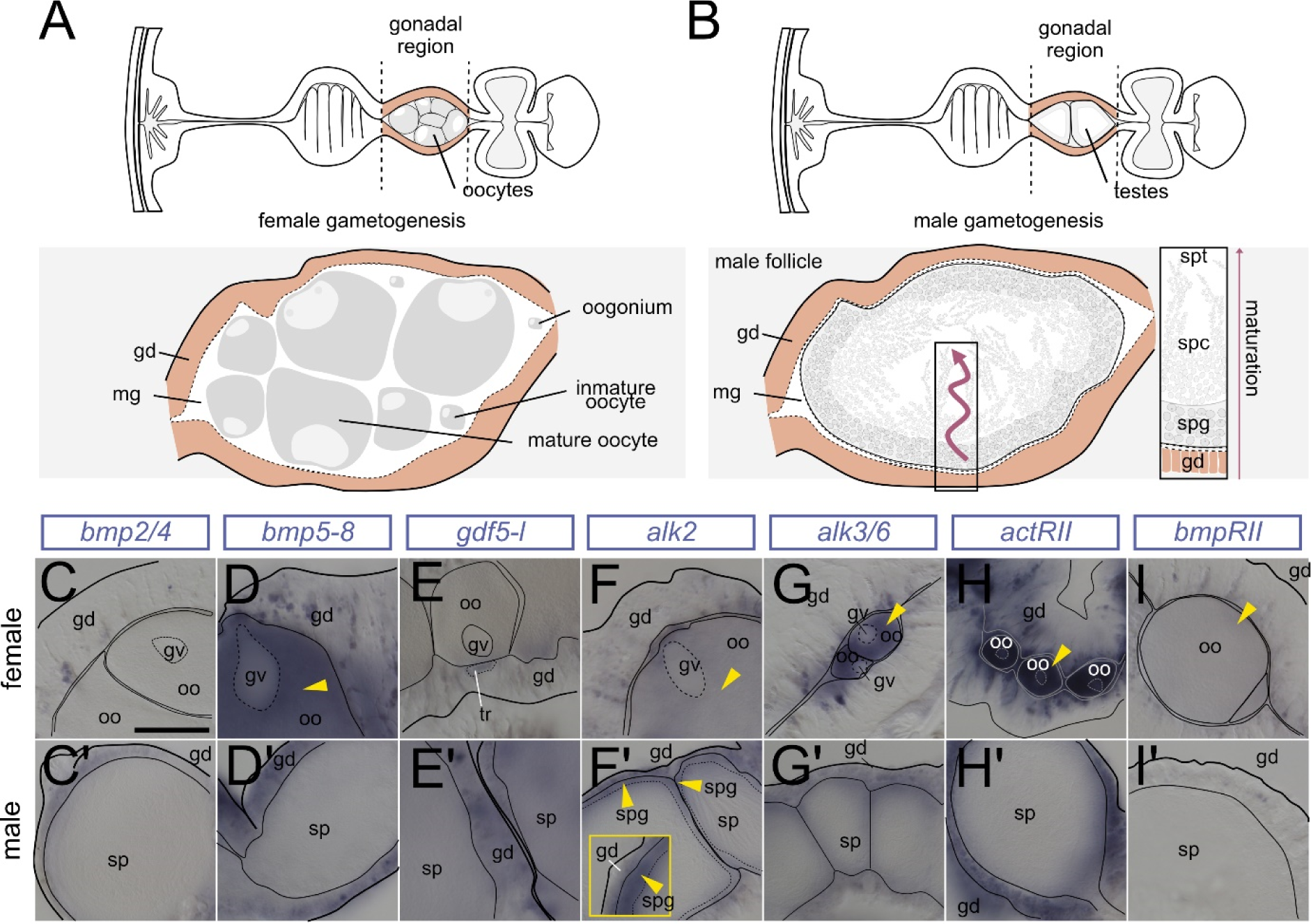
Expression of BMP ligand and receptor genes in maturing female and male gametes. (A-B) Schematic of the mesentery morphology at the level of the gonad as transversal cross-section, gonad region is highlighted in pink. (A) oogenesis (B) spermatogenesis. (C-I’) BMP pathway gene expression analysis by ISH on cross-sections of the female (C-I) and male (C’-I’) gonadal region. (C-I) yellow arrowheads indicate detectable transcript in the oocytes (F’) *alk2* expression in spermatogonia (yellow arrowheads). Scale bar 100 µm; mg -mesoglea, gd - gastrodermis, oo - oocyte, gv - germinal vesicle, sp - spermaries, spg – spermatogonia, spc – spermatocytes, spt – spermatids and sperm.

### BMP signaling is absent from vasa-positive germ cells and stem cells

Single-cell transcriptomic analyses suggested the upregulation of BMP pathway genes in the previously identified stem cell cluster (Cole et al., 2023; Cole et al., 2024; Steger et al., 2022). While the *Nematostella* stem cell system is still poorly characterized, vasa2 protein was reported to not only mark primordial germ cells, maturing oocytes and the spermatogonia ((Chen et al., 2020; Extavour et al., 2005; Praher et al., 2017), see **Figure 8A-B**) but also a recently described population of putative stem cells in the mesentery contributing to germline and somatic lineages (Miramón-Puértolas & Steinmetz, 2023). To determine if BMP signaling is active in any of these populations in the mesentery, we performed immunostaining against pSMAD1/5 in combination with the monoclonal antibody against *Nematostella* vasa2 (Praher et al., 2017). In agreement with previous studies, we detected vasa2-positive basiepithelial cells in the reticulate and gonad tracts in the subpharyngeal (**Figure 8C-C’**) and gonadal region (**Figure 8D-D’**), yet there was no overlap with the BMP signaling activity (**Figure 8C’’, 8D’’**). In rare instances, we detected pSMAD1/5-positive basiepithelial cells located in the gonad region (**Figure 8D’’,** blue arrowhead) that were, however, distinct from vasa-positive cells (**Figure 8D’’**, white hollow arrow). In line with what has previously been described by us and others, vasa2 protein was present in maturing oocytes and basiepithelial germ cells in the female gonad, and in the outermost region of the spermaries populated by the spermatogonia (Chen et al., 2020; Denner et al., 2023; Miramón-Puértolas & Steinmetz, 2023; Praher et al., 2017), yet gametes and putative germ cells displayed no pSMAD1/5 immunoreactivity (**Figure 8E-F’’**). Occasionally, we observed pSMAD1/5-positive cells that were negative for vasa2 protein in close proximity to the vasa-positive spermatogonia (**Figure 8F’-F’’’**).

**Figure 8.**
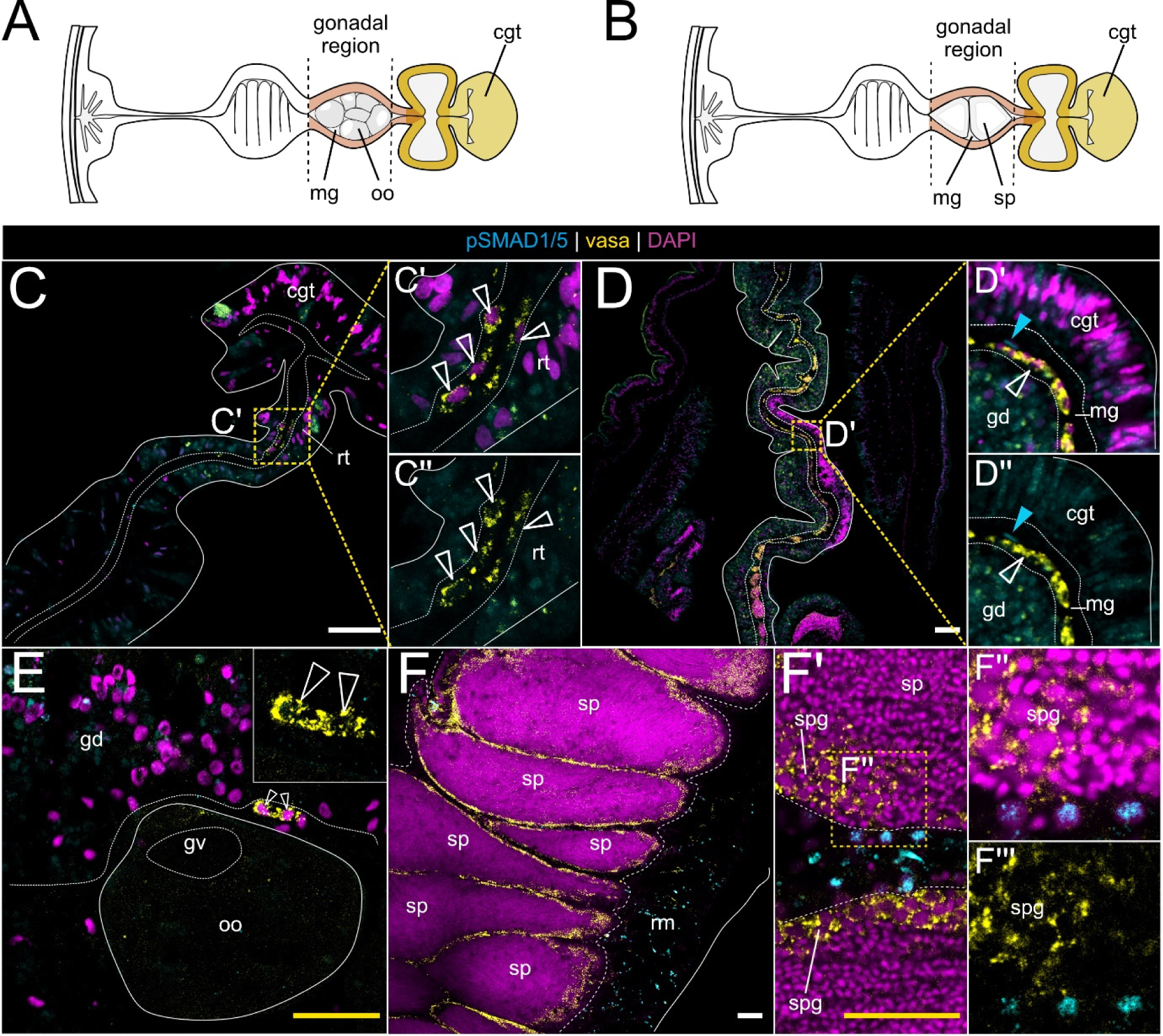
BMP signaling is absent from vasa2-positive germ and stem cells. (A-B) Schematic of the mesentery morphology at the level of the gonad region in female (A) and male (B) on a transverse section. Gonad highlighted in pink, septal filament highlighted in shades of yellow. (C-F) immunostaining of pSMAD1/5 (cyan), vasa (yellow) and DNA (magenta) in different parts of the gonad region. (C) vasa2-positive basiepithelial cells in the reticulate tract are pSMAD1/5-negative. (D) In the gonad region, vasa2-positive basiepithelial cells are pSMAD1/5-negative. Rarely, pSMAD1/5-positive, vasa2-negative basiepithelial cells are present. (E) vasa2-positive putative germ cells and maturing oocytes are pSMAD1/5-negative. (F) vasa2 accumulation in the spermatogonia. (F’-F’’’) sporadic pSMAD1/5-positive cells in the male gonad do not overlap with vasa2 in the spermatogonia. mg – mesoglea, oo - oocyte, sp – spermaries, cgt - cnidoglandular tract, rt - reticulate tract, rmr - retractor muscle region, gd - gastrodermis, gv - germinal vesicle, spg - spermatogonia.

In summary, we did not observe an overlap between vasa2-positive basiepithelial cells and pSMAD1/5-positive basiepithelial cells. Strikingly, BMP signaling is active in basiepithelial cells located in the basal mesentery region. These cells are vasa2-negative, however, it is conceivable that they may represent a progeny of vasa2-positive stem cells and cluster together with the stem cells in the single-cell transcriptomic data.

### BMP signaling in the gonad region is active in female trophonemata and male ciliated plugs

Another exception to the low BMP signaling activity of the medial mesentery we discovered in the accessory cells of the somatic gonad – the trophonema cells specific for the female, and ciliated plug cells specific for the male (Denner et al., 2023). They form small islands in the somatic gonad gastrodermis in close proximity to maturing gametes (**Figure 9A-B**). In the accessory cells of both, male and female polyps, we detected low levels of pSMAD1/5 activity (**Figure 9C-F**). In our in situ hybridization analysis of the somatic gonad, we could not observe detectable levels of BMP receptor expression, however, trophonema and ciliated plug cells were positive for the BMP ligand *gdf5-l* and the BMP antagonist *chordin* RNA (**Figure 9G-L**).

**Figure 9.**
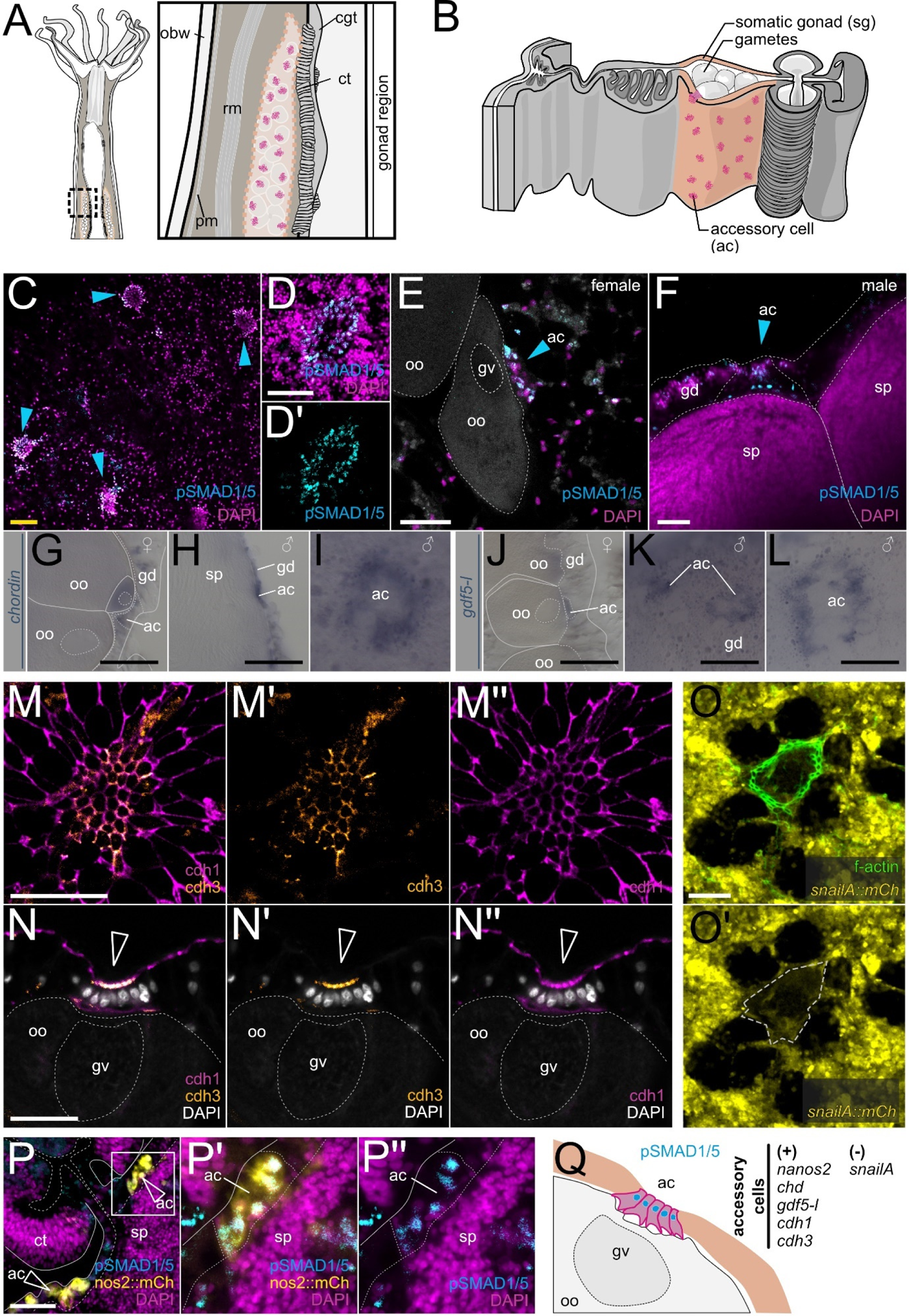
BMP signaling is active in accessory cells in the somatic gonad gastrodermis. (A-B) Schematics of the mesentery morphology in the gonad region in a lateral view (A) and in a transverse section (B). Gonadal region highlighted in light pink, accessory cells in magenta. (C-F) pSMAD1/5-positive accessory cells (blue arrowheads) in the gonad gastrodermis are in direct contact with (E) maturing oocytes and (F) spermaries. Expression of (G-I) *chordin* and (J-L) *gdf5-l* in female and male accessory cells. (M-N’’) cdh1 and cdh3 localization in gonad gastrodermis and accessory cells. cdh1 is present in the apical cell junctions throughout the gastrodermis, while cdh3 is specific to accessory cells (hollow white arrowheads on N-N’’). (O) f-actin staining shows apical constriction of accessory cells (O-O’) *snailA::mCherry* transgenic reporter (Kirillova et al., 2018) labels gastrodermal cells but it is excluded from accessory cells. (P-P’’) mCherry/pSMAD1/5 double-positive accessory cells in the *nanos2::mCherry* transgenic reporter line (Denner et al., accepted). Scale bars 100 µm (black), 50 µm (yellow), 25 µm (white). ac - accessory cells, sg - somatic gonad, gd - gastrodermis, oo - oocyte, gv - germinal vesicle, sp – spermaries, ct - ciliated tract;

While accessory cells have been addressed in histological and ultrastructure studies (Eckelbarger et al., 2008; Moiseeva et al., 2017), they could not be clearly assigned to any cell cluster by single-cell transcriptomic analyses. Since no distinct molecular signatures are known for the trophonemata and ciliated plug cells, we aimed to further analyze these cell populations. Trophonemata and ciliated plugs are apparent due to their tight aggregation and apical constriction of the cells in them compared to the circumjacent gastrodermal epithelium easily discernible in classical cadherin and fibrillar actin stainings (**Figure 9M-O’**). *Nematostella* has two classical cadherins, cdh1, which is expressed in the gastrodermis of the mesenteries, and cdh3 expressed in the epidermis and in the pharynx-derived tissue including the septal filament, ((Pukhlyakova et al., 2019), **Figure 9 – Figure Supplement 1**). Surprisingly, in the otherwise cdh1-positive/cdh3-negative gonadal epithelium, cdh3 was clearly detectable in the contacts between the apically constricted cells of the trophonemata and ciliated plugs (**Figure 9M-N’’**). Side views of the trophonema (**Figure 9N-N’’**) showed colocalization of cdh3 and cdh1 at the apical side of the cells facing the gastric cavity, while only cdh1 was present at the basal trophonema side facing the animal pole of the oocyte, where the germinal vesicle is located (**Figure 9N-N’’**). In line with the differential expression of cadherins, we observed the downregulation of mCherry expression in the trophonemata in the *snailA::mCherry* reporter line compared to the rest of the gonad gastrodermis (**Figure 9O-O’**), suggesting the repression of cdh3 in the mesentery gastrodermis by SnailA, similar to the situation reported in the embryo (Pukhlyakova et al., 2019). Moreover, recently, trophocytes have been shown to express the RNA-binding protein Nanos2 (Denner et al., 2023), which we have recently identified as a direct target of pSMAD1/5 during early development (Knabl et al., 2024). In line with that, we found pSMAD1/5 activity overlapping with the mCherry expression in the *nanos2::mCherry* transgenic reporter line (**Figure 9P-P’’**).

### The medusozoan species *Aurelia coerulea* and *Tripedalia cystophora* display broad BMP signaling activity across different body regions

The expression of several BMP components, but mostly of BMP ligands, has been previously analyzed in several members of Medusozoa, including *Hydra*, *Clytia*, *Podocoryne* and *Aurelia* (Hobmayer et al., 2001; Kraus et al., 2015; Reber-Muller et al., 2006; Reinhardt et al., 2004), yet nothing is known about the activity and localization of BMP signaling in jellyfish and hydroids. Therefore, we aimed to test the cross-reactivity of the pSMAD1/5 antibody in multiple medusozoan representatives. We failed to detect specific anti-pSMAD1/5 in any of the assayed hydrozoan species including *Clytia hemisphaerica, Hydra vulgaris*, *Hydractinia sp.* and *Ectopleura sp.* and a scyphozoan species, *Sanderia malayensis* (**Figure 10 - Figure Supplement 1**). In contrast, we successfully detected BMP signaling by pSMAD1/5 staining in the scyphozoan jellyfish *Aurelia coerulea* and the cubozoan box jellyfish *Tripedalia cystophora*. Both species feature a triphasic life cycle common for most Medusozoa, with a free-swimming larva, a sessile polyp, and a free-swimming medusa stage (**Figure 10A, E**).

**Figure 10.**
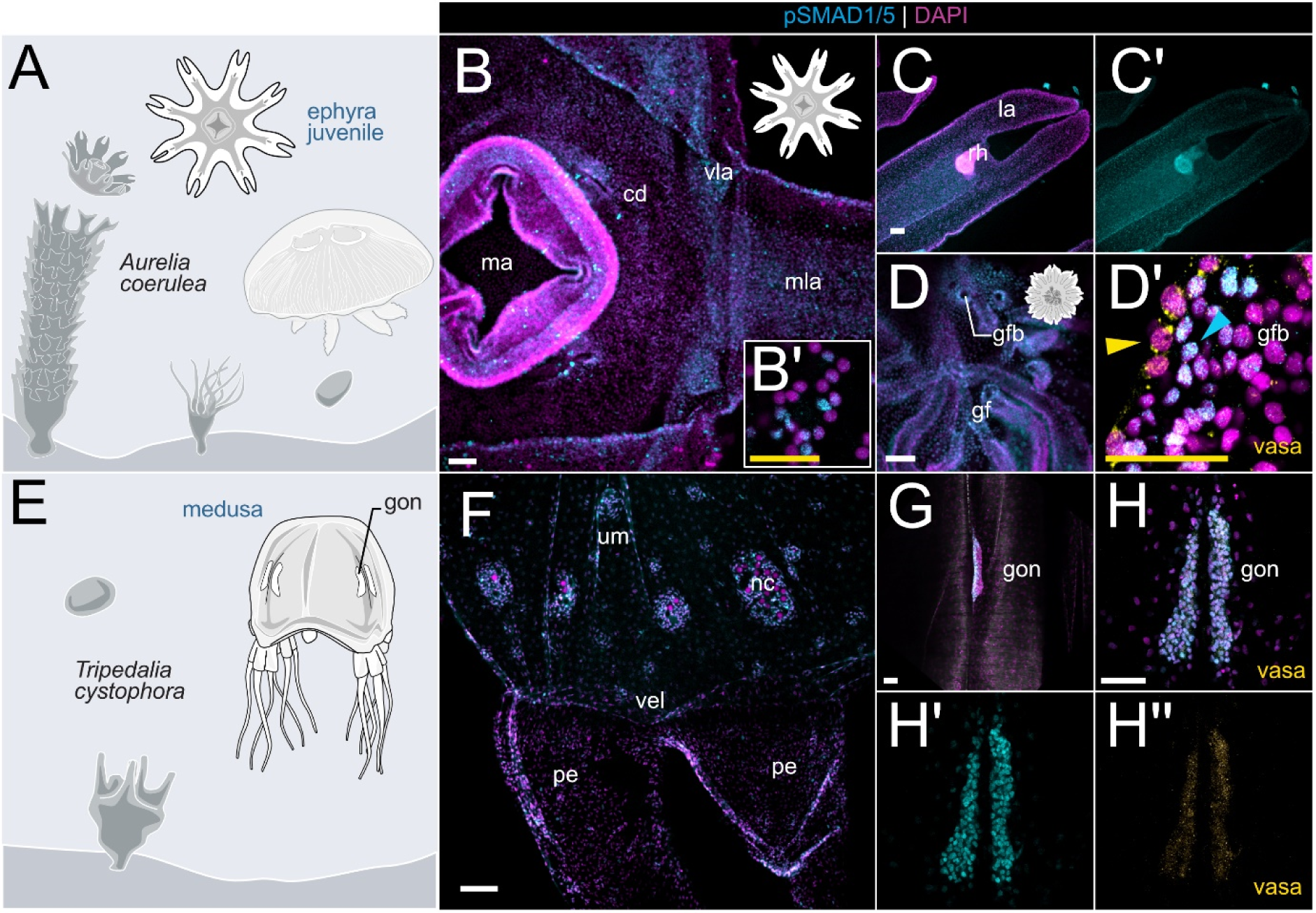
BMP signaling is broadly active in medusozoans, the scyphozoan jellyfish *Aurelia* and the cubozoan jellyfish *Tripedalia*. (A) Schematic life cycle of *Aurelia coerulea*. (B-D’) pSMAD1/5-positive nuclei in the central disc (B-B’), lappet and rhopalia (C-C’). (D-D’) vasa2-staining in the region of the gastric filaments (yellow arrow) shows no overlap with pSMAD1/5-positive nuclei (blue arrow). (E) Schematic life cycle of *Tripedalia cystophora*. (F) Broad pSMAD1/5 activity in the umbrella and velarium. (G) Tripedalia gonad is pSMAD1/5-positive. (H-H’’) Germ cells in the *Tripedalia* gonad are pSMAD1/5- and vasa-positive. Scale bar 50 µm (white) and 25 µm (B’, yellow). ma – manubrium, cd – circular disc, vla – velar lappet, mla -marginal lappet, gf – gastric filament, gfb – gastric filament base, um – umbrella, nc – nematocyst wart, vel – velarium, pe – pedalium, gon – gonad;

In *Aurelia*, we focused on the developmental stages that were most accessible for whole mount immunofluorescence staining: the polyp stage and early juvenile medusa stages (ephyra and metaephyra). *Aurelia* polyp (scyphistoma) undergoes serial transverse fission and releases multiple genetically identical juvenile jellyfish (ephyrae), which develop into the metaephyra and, subsequently, into adult medusae (**Figure 10A**). No active BMP signaling was observed in the unsegmented *Aurelia* polyp (**Figure 10 – Figure Supplement 1**), while at the ephyra stage, BMP signaling activity was present across different portions of the body. Here, mixed populations of pSMAD1/5-positive and -negative cells were found in the epithelium of the central disc, the lappets and the sensory organs called rhopalia (**Figure 10B-C**). In contrast, the manubrium was free of BMP signaling, although we observed unspecific (non-nuclear) staining of mucoglandular cells (**Figure 10B**), similar to nonspecific staining observed in the *Nematostella* pharynx and septal filament. At the metaephyra stage, characterized by the stepwise fusion of the individual lappets into a bell-shaped umbrella, BMP signaling was observed in similar areas as in the ephyra. In addition, pSMAD1/5-positive cells were found at the base of the gastric filaments (**Figure 10D**), which are located in four distinct gastric pouches around the manubrium. In contrast, gastric filaments, which are very rich in gland cells, showed only unspecific (non-nuclear) staining. Later during development, the gastric pouches also contain four horseshoe shaped gonads sitting underneath the gastric filaments. Co-immunostaining with anti-pSMAD1/5 and anti-*Nematostella* vasa2 antibody revealed vasa-positive cells with a typical perinuclear vasa signal at the base of the gastric filaments that were distinct from the pSMAD1/5-positive cells located immediately next to the vasa expressing cells – similar to the situation in *Nematostella* (**Figure 10D’**).

In *Tripedalia*, antibody staining revealed a broad pSMAD1/5 activity in juvenile and adult jellyfish. Mixed populations of pSMAD1/5-positive and -negative cells were detectable in the umbrella, velarium, ring nerve, parts of the nematocyst warts and rhopalia of the jellyfish, while the activity was absent from the manubrium, pedalia and tentacles (**Figure 10F**). In sexually mature females, four pairs of wing-shaped gonads are located on the inside of the medusa bell next to the radial canals of the gastrovascular system. Strikingly, germ cells in the female gonad of *Tripedalia* were pSMAD1/5-positive (**Figure 10G-H’**), and co-staining with anti-*Nematostella* vasa2 antibody showed an overlap with the accumulation of vasa protein (**Figure 10H-H’’**), which was never observed for germ cells in *Nematostella*.

## Discussion

### Diversity of BMP signaling domains in Cnidaria indicates a variety of functions

To date, most of our knowledge on BMP signaling in Cnidaria comes from expression-based studies in Medusozoa, as well as expression-based and functional studies in anthozoan embryos (Finnerty et al., 2004; Genikhovich et al., 2015; Knabl et al., 2024; Kraus et al., 2015; Leclere & Rentzsch, 2014; Matus, Pang, et al., 2006; Matus, Thomsen, et al., 2006; Reber-Muller et al., 2006; Reinhardt et al., 2004; Rentzsch et al., 2006; Rentzsch et al., 2007; Saina et al., 2009). Research in anthozoans, however, only addressed the role of BMP signaling in directive axis establishment and patterning. In this work, we aimed to expand this understanding by investigating the activity of BMP signaling unrelated to axial patterning in the adult polyp of the anthozoan *Nematostella,* and two radially symmetric medusozoans, the juvenile ephyra of the scyphozoan *Aurelia* and the cubozoan jellyfish *Tripedalia,* utilizing an antibody against the BMP effector pSMAD1/5 (Genikhovich et al., 2015; Knabl et al., 2024; Leclere & Rentzsch, 2014). In all three species, BMP signaling was active in distinct cell populations in different locations, suggesting that BMP signaling must be involved in the regulation of multiple processes.

We centered our analysis on *Nematostella* and systematically dissected BMP activity domains in this bilaterally symmetric model sea anemone. We showed that BMP signaling was strongest in distinct areas of the epidermis of the oral region, as well as in specific cell populations in the gastrodermis of the mesenteric folds (**Figure 11A-D**). In the mesentery, pronounced BMP activity was detected in the basally located neuro-muscular region, while it appeared reduced, although not absent, in the more apical parts including the gonad region. In line with that, our analysis of the published single-cell transcriptomic data indicates broad expression of the BMP pathway genes (Cole et al., 2023; Cole et al., 2024; Steger et al., 2022). Expression of BMP receptor genes *alk2*, *alk3/6*, *bmprII* and *actrII* and BMP effector genes *smad1/5* and *smad4* was detectable in multiple cell clusters including “mat.cnido”, “retractor muscle”, “gastrodermis”, “epithelia.ecto”, “ectoderm.embryonic”, “pSC”, and “neuronal” cells. The diversification of BMP activity in multiple specialized cell types may already start during larval development, after patterning of the directive axis is completed, as indicated by our previous anti-pSMAD1/5 ChIP analysis. We identified direct BMP signaling target genes at late gastrula, when the axis patterning gradient of BMP signaling activity has just formed, and in 4d planula, when the axis is already patterned, and the pSMAD1/5 gradient is no longer present. While at both stages, many BMP targets are developmental regulators involved in the patterning of the directive axis, in 4d planula we observe an increasing number of targets with other putative functions such as metabolism, extracellular matrix dynamics or neurogenesis (Knabl et al., 2024). In the future, it will be important to follow the dynamics of BMP signaling between 4d planula and adult and figure out how adult BMP signaling domains appear.

**Figure 11.**
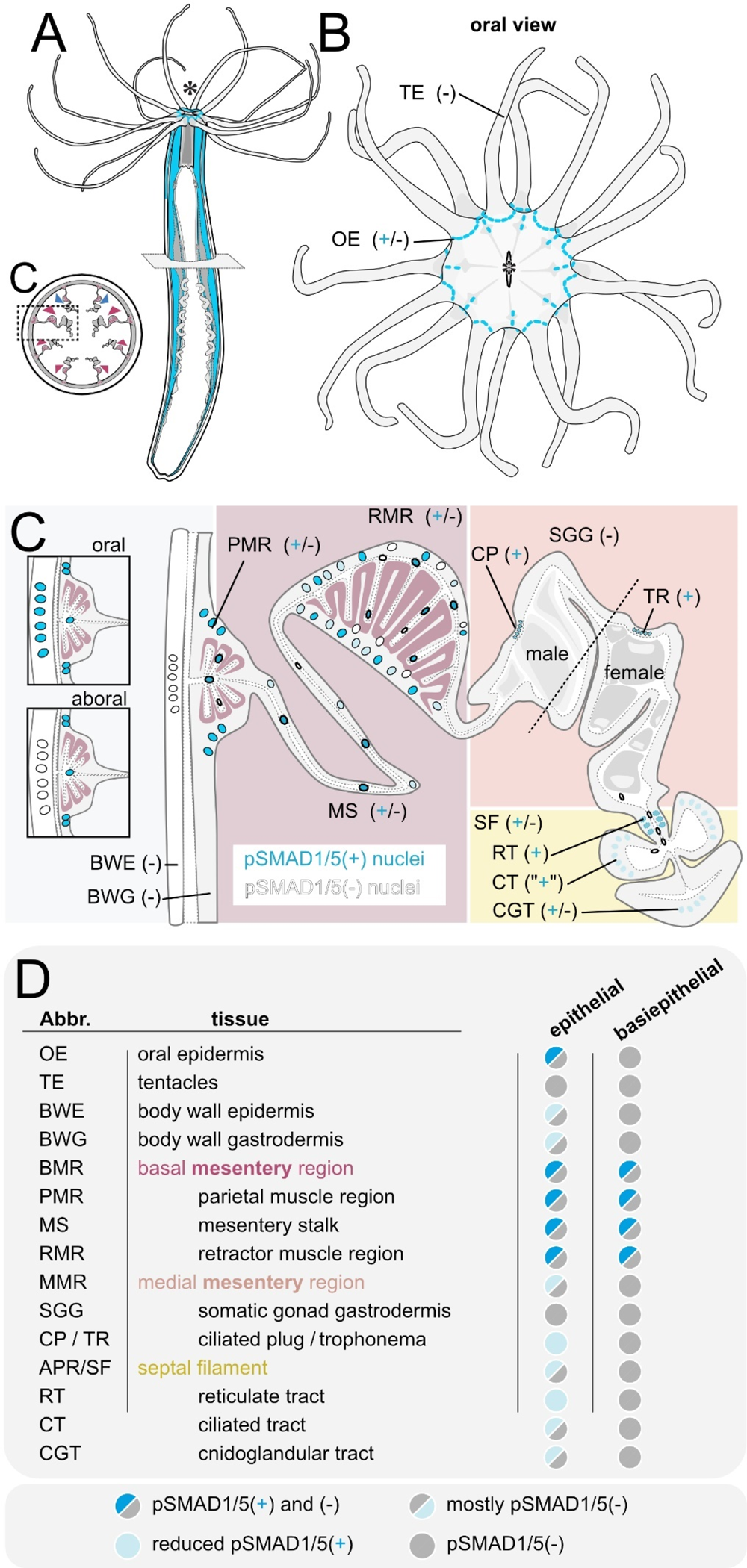
Summary of BMP signaling domains in the adult *Nematostella* polyp.

In medusozoans, BMP activity and the expression of BMP components appears to be similarly versatile. Our study revealed pSMAD1/5 being broadly active in the epithelium of the central disc, radial canal and lappets, in the rhopalia, as well as in the neurons and cnidocytes of *Aurelia* and *Tripedalia*. In line with that, earlier studies in the ephyra of *Aurelia* and medusa of *Clytia* and *Podocoryne* detected *bmp5/8* expression in the radial canal and sensory structures (rhopalia) (Kraus et al., 2015). In *Hydra, HySmad1* is absent from the tentacles and the peduncle but broadly detectable along the entire body column, where it is expressed in a variety of cell types including epithelial cells, neurons, gland cells, interstitial stem cells and endodermal epithelium as determined by northern blot (Hobmayer et al., 2001). In contrast, a *bmp2/4*-like gene and three copies of *bmp5-8* are expressed at the tentacle base, as well as in the body column (Krishnapati et al., 2020; Reinhardt et al., 2004; Rentzsch et al., 2007; Siebert et al., 2019; Watanabe, Schmidt, et al., 2014; Wenger et al., 2019)

Together, our data on BMP activity domains in three cnidarian representatives, combined with existing cnidarian expression data, indicate that BMP signaling may be relevant in at least three different contexts, besides axial patterning: 1) in tentacle formation, 2) in the neuro-muscular region, and 3) in the reproductive region.

### Potential role of BMP signaling in tentacle formation

Among the most pronounced domains of BMP signaling was the pSMAD1/5 activity between the tentacles (**Figure 11B, D**). A similar pattern of BMP activity can be observed during postembryonic development, when BMP signaling forms a cross-like activity domain in the oral epidermis ((Knabl et al., 2024), **Figure 2 – Figure Supplement 1A-A’**), with four pSMAD1/5-free territories at the localization of the four primary tentacle buds. Starting from the 4d planula, pSMAD1/5 activity is also accompanied by the expression of several BMP components and BMP target genes in the tentacle area. While pSMAD1/5-negative tentacle buds express genes encoding BMP2/4, BMP5-8 (**Figure 2 - Figure Supplement 1B-C**), the BMP antagonists Noggin, Gremlin (Matus, Pang, et al., 2006), the metalloprotease Tld (Matus, Thomsen, et al., 2006) and the BMP and BMP receptor binding molecule RGM (**Figure 2 - Figure Supplement 1D-D’,** (Leclere & Rentzsch, 2014)), the pSMAD1/5-positive intertentacular area expresses genes encoding a BMP binding protein CV2 ((Mörsdorf et al., 2024), **Figure 2 - Figure Supplement 1E-E’**) and another BMP ligand ADMP (**Figure 2 - Figure Supplement 1F-F’**). Single-cell transcriptomic data indicates that *admp* is the only BMP gene, whose expression persists in the epidermis of the adult polyp making it the prime candidate for signaling in the intertentacular domain at this stage.

Besides the expression of BMP signaling components, several other observations suggest the importance of BMP signaling in the tentacle region. Knockdowns of BMP signaling components *bmp2/4*, *bmp5-8*, *chd* and *rgm* not only interfere with the formation of the directive body axis and of the mesenteries (Genikhovich et al., 2015; He et al., 2018; Leclere & Rentzsch, 2014; Saina et al., 2009), but also cause a late phenotype at the stage corresponding to the primary polyp in controls. Knockdown animals become “worm-like”, displaying severe body elongation, everted pharynx and, importantly, lacking tentacles (He et al., 2018; Leclere & Rentzsch, 2014; Stokkermans et al., 2022). Another phenotype associated with tentacle formation was observed when we generated *bmp2/4* knockout animals using CRISPR/Cas9 (**Figure 11 – Figure Supplement 1A-B**). While most injected F0 embryos developed a “worm phenotype” and died, some embryos survived until adulthood. Many of these adults formed large bubble-like structures on the sides of their bodies. Genotyping showed that in the F0 animals, the bubbles contained *bmp2/4* mutant cells, and once we obtained heterozygous *bmp2/4^+/-^* F1, these animals also started sporadically developing bubbles (**Figure 11 – Suppl. 1B-C**). Morphological analysis confirmed the presence of epidermal muscles and spirocytes in the bubble tissue (**Figure 11 – Suppl. 1D-E’**). Since epidermal muscles and spirocytes (a specific cnidocyte type) are unique features of a normal *Nematostella* tentacle not present anywhere else in the animal, we speculate that bubble formation is a result of the ectopic and incomplete activation of the tentacle formation program.

Expression data suggest the association of BMPs with the tentacles in several Medusozoa. In the polyps of the hydrozoans *Hydra*, *Podocoryne* and *Clytia*, as well as in the polyp of the scyphozoan *Aurelia,* BMP ligand *bmp5-8* is expressed in developing tentacle buds (Kraus et al., 2015; Reber-Muller et al., 2006; Reinhardt et al., 2004). In the case of *Hydra*, several other BMPs, including *bmp2/4-like* and *bmp5-8c*, also show increased expression at the tentacle base in a homeostatic polyp, and all *bmp* genes react dynamically to budding and/or head regeneration (Reinhardt et al., 2004; Watanabe, Schmidt, et al., 2014). Our preliminary results indicate that treatment with a pharmacological BMP receptor inhibitor K02288 supports a role of BMP signaling in tentacle development. Like *Hydra*, *Nematostella* polyps have a high regenerative potential and, after bisection across the OA axis, both halves can regenerate the missing oral or aboral structures (Bossert et al., 2013; DuBuc et al., 2014; Passamaneck & Martindale, 2012). We observed that *Nematostella* polyps failed to re-establish their tentacles after head amputation when incubated in the K02288 inhibitor (10/10). Similarly, no tentacles were formed in *Hydra* polyps treated with K02288 after head amputation (10/10).

Taken together, we see evidence for the involvement of BMP signaling in regulating patterning and morphogenesis of the tentacle domain in *Nematostella* as well as in several medusozoans. However, the exact role of BMP signaling in this and the way BMP signaling acts in concert with other pathways, e.g. with FGF and Hedgehog signaling in *Nematostella* (Chen et al., 2020; Ikmi et al., 2020), remain to be discovered.

### Potential role of BMP signaling in neurons, muscles and cnidocytes

Our analysis of the single-cell transcriptomic data shows upregulation of BMP pathway genes in *Nematostella* neurons, muscles and cnidocytes. Neurons and cnidocytes are generated by a common pool of SoxB2-positive multipotent progenitors during embryonic, larval, and post-larval development (Richards & Rentzsch, 2014; Steger et al., 2022), while muscles originate from other cells. Our data in the adult polyp demonstrate that domains of pronounced BMP signaling activity coincide with the basal part of the mesentery (**Figure 11C-D**), and include two of the three neuron-rich areas of the gastrodermis: the longitudinal nerve tracts at base of the mesentery located next to the parietal muscle, and the intramesenterial neurons, some of which are embedded into the retractor muscle, but not the neurons of the body wall and the circular muscles.

ChIP-seq with an anti-pSMAD1/5 antibody on late gastrula and 4d planula of *Nematostella* revealed multiple genes as BMP targets including *arp6*, *ashB, atoh7*, *hmx3*, *isl*, *irx*, *lmx*, *soxC, rgm*, *ephrinB* and *netrin*, which are associated with neural function in vertebrates and invertebrates (Knabl et al., 2024). In contrast, we do not find many muscle-related genes among the direct BMP signaling targets, which suggests mesenterial neurons rather than parietal or retractor muscle cells as more likely candidates for being pSMAD1/5-positive. Despite being vital during formation and patterning of the centralized nervous system in many Bilateria, the roles of BMP signaling in the formation of the diffuse nerve net common in Cnidaria are largely unknown. In *Nematostella,* early neurogenesis is considered to be independent of BMP signaling (Rentzsch et al., 2017; Watanabe, Kuhn, et al., 2014), with pSMAD1/5 becoming detectable in the oral domain during gastrulation, which is after the onset of the neural gene expression at the blastula stage and in a different location in comparison to where the first RF-amide-positive and elav-positive neurons are found in the gastrula (Knabl et al., 2024; Nakanishi et al., 2012; Richards & Rentzsch, 2014). In line with that, knockdown and upregulation of BMP signaling showed no effects on early neurogenesis, however, an anti-as well as a pro-neurogenic potency of BMP signaling has been observed during oral nerve net formation in the planula larvae (Watanabe, Kuhn, et al., 2014).

While some neurons appear to be pSMAD1/5-positive, the situation with cnidocytes is different. Transcripts of the BMP pathway genes are expressed in mature cnidocytes in the single-cell data (**Figure 1E**), however, we did not observe nuclear pSMAD1/5 in any stinging cell subtype, besides the unspecific, non-nuclear signal in spirocytes (**Figure 2 – Figure Supplement 2**). Notably, the cnidocyte-specific expression of some of the BMP components may be shared with other cnidarians, given the fact that HySmad1 could be detected in mature nematocytes of the gastric region in *Hydra* (Hobmayer et al., 2001). Co-staining of pSMAD1/5 in different existing transgenic reporter lines (Renfer et al., 2010; Richards & Rentzsch, 2014; Steger et al., 2022) or together with antibody staining for neuronal, muscle, and nematocyte markers will help elucidate the role of BMP signals in the development, regional patterning, and maintenance of neurons, cnidocytes and muscles in *Nematostella* and, eventually, other cnidarians.

### Potential role of BMP signaling in the reproductive region

While we have a comprehensive understanding of the stem cell system in hydrozoan species (Houliston et al., 2010; Seipel et al., 2004; Siebert et al., 2019; Varley et al., 2023), most efforts to characterize stem cells in non-hydrozoan cnidarians remained unsuccessful. Ultrastructural studies in Anthozoa and Scyphozoa revealed extraepithelial, amoeboid cells, which were proposed to be stem cells (Gold & Jacobs, 2013; Tucker et al., 2011), but lack further characterization. A recent study in *Nematostella* provided evidence for a *vasa2*/*piwi1* double-positive basiepithelial putative stem cell population in the mesentery in the basal portion of the septal filament (Miramón-Puértolas & Steinmetz, 2023). As these adult stem cells contribute to vasa2-positive primordial germ cells (PGCs) as well as to the soxB2-positive neuronal progenitors, they were proposed to be homologous to the interstitial stem cells (i-cells) of Hydrozoa (Miramón-Puértolas & Steinmetz, 2023).

Our analysis of *Nematostella* scRNA-seq data indicated the conspicuous upregulation of BMP pathway members in the previously identified stem cell cluster (Cole et al., 2023; Cole et al., 2024; Steger et al., 2022). In accordance with the observations by Miramon et al., 2023, we detected vasa2-positive basiepithelial cells in the septal filament (**Figure 8C-D**), however, these cells were pSMAD1/5-negative. Instead, we found upregulated expression of BMP ligands *bmp2/4* and *bmp5-*8 as well as pSMAD1/5-positive cells in the epithelial parts of the septal filament, where vasa-positive basiepithelial cells concentrate. Curiously, pSMAD1/5 is detectable in vasa2-negative basiepithelial cells located in the neuromuscular domain of the mesentery. Considering that single-cell transcriptomic data indicated the upregulation of BMP pathway genes in putative stem cells, these findings suggest that either *vasa2*/*piwi1*-positive adult stem cells express BMP components but do not receive BMP signals or that there are other, *vasa*-negative stem cell populations that may be pSMAD1/5-positive.

Curiously, expression data of BMP components in Hydrozoa also implies possible involvement of BMP signaling in interstitial stem cell and germ cell development. Hydrozoan i-cells constitute a pool of pluri-or multipotent stem cells that contributes to somatic cells and the germline (Bosch & David, 1987; Holstein, 2023; Houliston et al., 2010; Muller et al., 2004; Varley et al., 2023). In Hydra, Northern hybridization of *smad1/5* indicated that it is upregulated in i-cells (Hobmayer et al., 2001), while transcriptomic analysis showed the upregulation of the BMP receptor *actr1* in multipotent interstitial cells and *smad1/5* and BMP antagonist *dand* in germline stem cells (Nishimiya-Fujisawa et al., 2023). In *Hydractinia*, the expression a putative *bmp receptor* has been detected in developing oocytes and the male gonophore gastrodermis (Sanders et al., 2014). *bmp5-8* has been also found to be differentially regulated in female and male *tfap2*-positive germ cells in *Hydractinia* (DuBuc et al., 2020). Another TGFβ molecule, *gonadless* (*Gls*), which may or may not be a BMP, is expressed in *tfap2*-positive germ cells, and *gls* knockout does not affect the germ cells but results in the absence of the sexual zooids carrying gonads in *Hydractinia* (Curantz et al., 2024).

All these observations in hydrozoans imply a role of BMP signaling in gonado- or gametogenesis. This role appears to be highly conserved across Bilateria (Lochab & Extavour, 2017), and, at least in part, is supported by our study. Single-cell transcriptomic data from the adult *Nematostella* polyp suggest that BMP genes are not expressed in primordial germ cell (PGC) clusters, and we never observed pSMAD1/5 activity in the vasa2-positive stem and germ cell lineage, including maturing gametes (**Figure 8C-D**), however, RNA of several BMP signaling components was detected in the developing oocytes (*bmp5-8* and *alk6*) and spermatogonia (*alk2*). Similar to *Nematostella*, the juvenile *Aurelia* metaephyra displayed pSMAD1/5 positive cells in proximity to the developing reproductive region in the gastric pouches, but without showing an overlap with vasa-positive cells. Notably different is the situation in *Tripedalia*, where BMP signaling was specifically active in vasa-positive germline cells in the gonad.

One interesting cell population in this context are the cnidarian-specific accessory cells described in Anthozoa and Scyphozoa (Denner et al., 2023; Eckelbarger et al., 2008; Eckelbarger & Larson, 1992; Lauretta et al., 2018; Lebouvier et al., 2022; Moiseeva et al., 2017; Scott & Harrison, 2009; Tan et al., 2020; Wedi & Dunn, 1983). *Nematostella* accessory cells, termed trophocytes in the female and ciliated plug cells in the male polyp, are located in the somatic gonad in a close contact with maturing oocytes and the spermatogonia (**Figure 9C-F**). Little is known about their function, however, unlike previously suggested, trophonemata appear not to be involved in nutritional transport (Lebouvier et al., 2022) but might play a role during germ cell development (Denner et al., 2023). While the somatic gonad gastrodermis is generally pSMAD1/5-negative, we detected active BMP signaling in accessory cells of the female and male polyp, also expressing the BMP ligand *gdf5-l* and the BMP antagonist *chordin*. Accessory cells display an upregulation of RNA-binding zinc finger molecule *nanos2*, a gene expressed in the piwi1/vasa2/nanos2-positive multipotent stem cell population and in a broad progeny of the neuroglandular lineage (Denner et al., 2023). Furthermore, *nanos2*^-/-^ knockouts not only lack accessory cells but also fail to establish the germline (Denner et al., 2023). While we cannot comment on the regulatory relationship of BMP signaling and *nanos2* expression in the adult polyp besides them being active together in accessory cells, *nanos2* is as a direct target gene of pSMAD1/5 during larval development (Knabl et al., 2024).

## Conclusion

In this study, we provide a whole-body atlas of BMP signaling activity in the adult bilaterally symmetric cnidarian model *Nematostella* and a brief overview of BMP signaling activity in two radially symmetric cnidarian models, *Aurelia* and *Tripedalia*. Currently, we are unable to state with certainty which cell types/states from single-cell transcriptomic data are the ones receiving BMP signals in the adult *Nematostella* polyp, however, in the future, as more and more adult cell states become “de-orphanized”, this question will no doubt be resolved. The same approach will then have to be expanded to our two medusozoan models, *Aurelia* and *Tripedalia*. Our analyses show that BMP signaling is involved in a variety of processes beyond axial patterning, which explains the presence of the BMP signaling components in non-bilateral Cnidaria. Among these processes, the role of BMP signaling in regulating tentacle formation, neuronal differentiation and gameto- or gonadogenesis clearly stand out and will form focal points of the future studies.

## Methods

### Animal culture

Nematostella vectensis adult polyps, separated by sexes, were maintained in the dark at 18°C in 16 ppm artificial seawater (*Nematostella* medium, NM) and spawned as described before (Fritzenwanker & Technau, 2002; Genikhovich & Technau, 2009). To generate BMP2/4 mutants, the genomic sequence encoding for the N-terminus of the mature BMP2/4 ligand was targeted using the ALT-R CRISPR-Cas9 system (IDT). crRNAs targeting two (overlapping) sequences were ordered: AAACGGAGCCTGCGGTCCGGCGG and AGAAAACGGAGCCTGCGGTCCGG, and both proved to work efficiently. Genotyping was performed using the primers GCAGACAACGATGGGATTGACGCTAGTG and GTATTGCCCGTTCTAATCATCCTTGAAGGC.

### Fixation for immunohistochemistry

Adult animals were starved for up to 3 days and left in 0.1M MgCl2 in NM until fully relaxed (5-15 min). For fixation, polyps were transferred to a new dish containing ice-cold fixation solution 1 (3,7% Formaldehyde in PBS). For antibody stainings of pSMAD1/5, fixation solution 1 (3.7% Formaldehyde, 1x PBS, 0.2% Triton-X) was used. For antibody stainings of pSMAD1/5, vasa2, cdh1 and cdh3, fixation solution 2 (3.7% Formaldehyde, 0.4% glyoxal, 0.1% MeOH, 1x PBS, 0.2% Triton-X) was used. To ensure consistent penetration, the fixation solution was injected slowly through the mouth opening, using a 1 ml syringe and a Sterican blunt needle with a bent tip. Polyps were transferred to a new tube with fresh, ice-cold fixation solution and fixed for 60 min under rotation at 4°C. After 20-30 min fixation time, samples were transferred to a dish with fresh fixation solution for further dissection. For vibratome sectioning, samples were cut into ∼5 mm pieces transversely through the body column samples. For whole mount imaging, the body column was opened longitudinally, and mesenteries were extracted from the body wall. After dissection, the fixation was continued under rotation at 4°C. The fixative was removed in 3 washes with 1x PBS / 0.2% Triton-X. To remove pigments, samples were washed with 100% methanol and repeatedly exchanged until the supernatant remained clear. Samples were washed 3x with 1xPBS, 0.2% Triton-X and directly further processed. For anti-pSMAD1/5 stainings, samples were incubated in Bloxall™Blocking Solution (Vector Laboratories) before the antibody blocking step (see Immunohistochemistry) to reduce unspecific signal. Samples were incubate with Bloxall for 10-30 min at room temperature, followed by 3x washes with 1x PBS.

### Fixation for in situ hybridization

Sample preparation for in situ hybridization of mesentery pieces or pieces of the body column was carried out as described before (see Fixation for Immunohistochemistry). Sample preparation and fixation was performed at room temperature using fixation solution 1 (3.7% FA, 1x PBS, 0.2% Triton-X), fixation was carried out on a rotator for 1 hour at room temperature. ISH samples were stored in 100% MeOH at -20°C.

### Immunohistochemistry

Samples were blocked in Blocking solution I (1% BSA, 5% sheep serum, 1x PBS, 0.2% Triton-X, 20% DMSO) for at least two hours at room temperature. In parallel, the primary antibodies were pre-absorbed in Blocking solution II (1% BSA, 5% SS, 1xPBS, 0.2% Triton-X, 0.1% DMSO). Primary antibody incubation of the sample was carried out overnight at 4°C with the following primary antibody dilutions: rabbit anti-pSMAD1/5/9 1:200 (mAb #13820, Cell signaling, (Genikhovich et al., 2015; Knabl et al., 2024; Leclere & Rentzsch, 2014), mouse anti-vasa2 1:1000 (Praher et al., 2017), rabbit anti-cadherin1 (1:500) and mouse anti-cadherin3 (1:1000) (Pukhlyakova et al., 2019), mouse anti-Tyrosinated Tubulin (TUB-1A2, T9028 Sigma, 1:1000). Samples were washed 10 times with 1x PBS / 0.2% Triton-X, blocked in Blocking solution II for 1 hour at room temperature and incubated with the secondary antibodies 1:1000 (goat α-rabbit IgG-Alexa568 (Invitrogen A11008), goat α-rabbit IgG-Alexa633 (Invitrogen A21070), goat α-mouse IgG-Alexa488 (Invitrogen A11001), goat α-mouse IgG-Alexa568 (Invitrogen A11004) and DAPI, 1:1000) in Blocking solution II at room temperature for two hours or at 4°C overnight. After 10 washes in 1x PBS / 0.2% Triton-X, were either infiltrated with and mounted in VECTASHIELD® Antifade Mounting Medium (H-1000-10, VectorLabs) or further processed for vibratome sectioning.

### Vibratome sectioning

Vibratome sectioning was performed after antibody staining or in situ hybridization. Embedding medium (10% gelatin in PBS) was dissolved at 50°C and stored at -20°C in 2 ml tubes. Before use, the embedding medium was warmed at 37°C until liquefied and pieces of the body column were left for infiltration until settling down to the bottom of the tube. Samples in the embedding medium were poured into mounting molds (cut 1 ml pipette tip), pieces were orientated, and molds were transferred to the fridge for hardening for 30 min. Hardened gelatine blocks were fixed in 4% Formaldehyde in PBS at 4°C overnight. Samples were washed 3x with 1xPBS and sectioned at a Leica VT1200 vibratome (speed = 0.5, amplitude= 0.7 and auto feed thickness = 50-100 μm;). Sectioned ISH samples were infiltrated with and mounted in glycerol, IHC samples in VECTASHIELD® Antifade Mounting Medium (H-1000-10, VectorLabs).

### in situ hybridization (ISH)

Urea-based in situ hybridization on adult tissues was carried out as described previously (Lebouvier et al., 2022), with minor changes. Pieces of Nematostella body column or of individual mesenteries were dissected and fixed for one hour at room temperature (as described in Fixation for immunohistochemistry and Fixation for in situ hybridization). Anti-Digoxigenin-AP (anti-DIG/AP) Fab fragments (Roche) were diluted 1:4000 in 0.5% blocking reagent (Roche) in 1x Blocking buffer solution (Roche).

### Single cell transcriptomic analysis

Single-cell transcriptomic analysis of BMP pathway genes was performed using previously published data by Steger et al. 2022, Cole et al. 2023 and Cole et al. 2024. The R-package Seurat vs4 (Stuart et al., 2019) was used to process the count matrices to include adult wild type libraries. The full dataset from Cole et al. 2024 was partitioned into samples derived only from adult tissues, and all clusters with less than 10 remaining cells were dropped. Expression of genes of interest across the available clustering was examined using the Seurat::DotPlot function, holding maximum dot size fixed at 100%.

## Acknowledgements

We thank Alison Cole for advice and feedback on the single-cell transcriptomic data analysis. This research was funded in whole or in part by the Austrian Science Fund (FWF) grant (DOI 10.55776/P32705) to G.G.. P.K. is a recipient of the Dimitrov Fellowship of the Austrian Academy of Sciences (OeAW). D.M. is a recipient of the Austrian Science Fund (FWF) Lise-Meitner Fellowship (DOI 10.55776/M3291). For the purpose of Open Access, the author has applied a CC BY public copyright license to any Author Accepted Manuscript (AAM) version arising from this submission. Confocal microscopy was performed at the Core Facility Cell Imaging and Ultrastructure Research, University of Vienna - member of the Vienna Life-Science Instruments (VLSI).

## Author contributions

P.K. designed and performed most of the experiments. D.M. generated and analyzed the *bmp2/4* knockout. G.G. conceived the project, acquired funding and designed experiments. P.K. and G.G. wrote the paper. All authors edited the paper.

## Data availability

All data necessary to evaluate the findings are presented in the main text or the supplementary materials.

---

**Figure 1 – Supplement 1.**
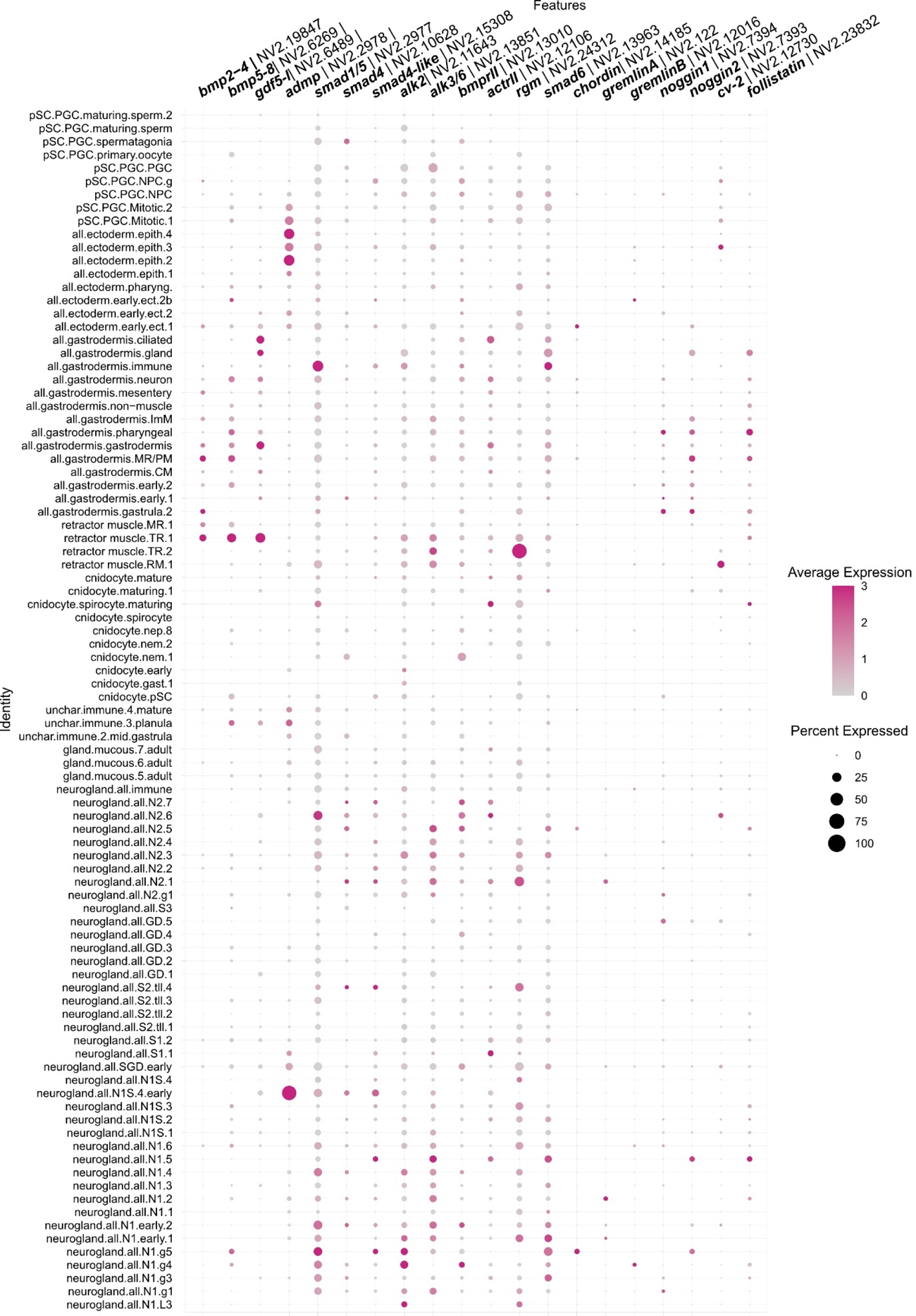
Dot plot showing the expression of BMP pathway genes in single-cell transcriptomic data of adult *Nematostella* polyp tissues with fine clustering presenting 91 transcriptomic cell states (Cole et al., 2023; Cole et al., 2024; Steger et al., 2022).

**Figure 2 – Figure Supplement 1.**
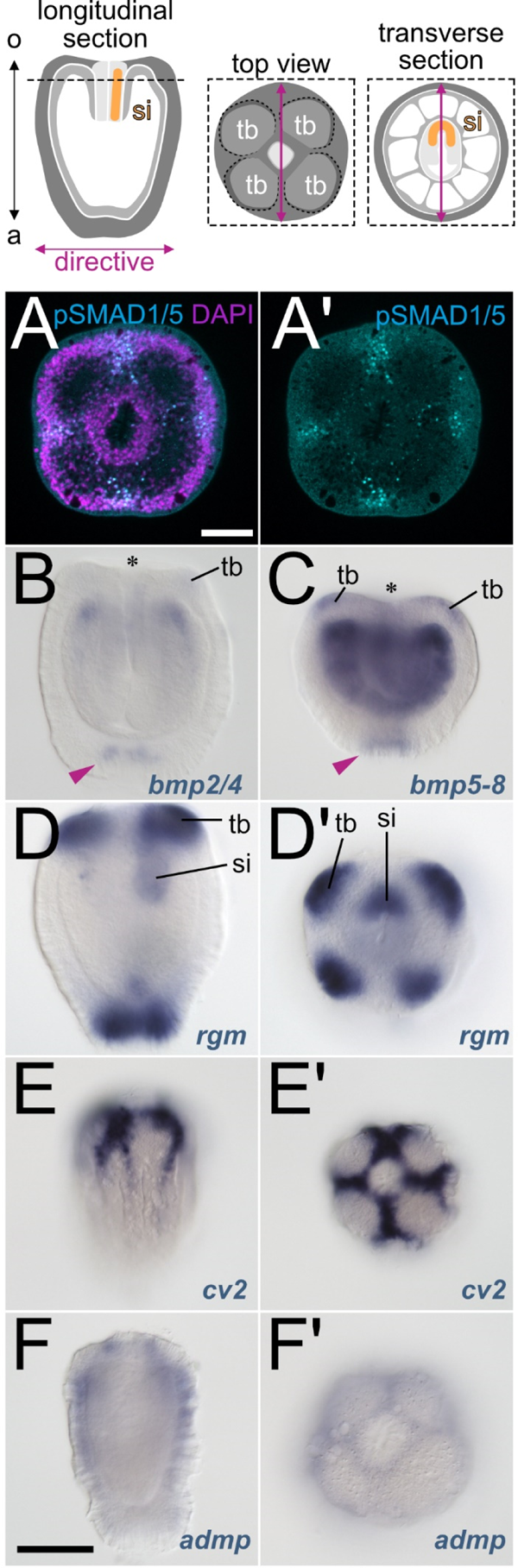
BMP signaling activity and expression of BMP signaling components in the 4d planula larva. (A-A’) pSMAD1/5 activity between the forming tentacles and around the mouth of the 4d planula. (B) *bmp2/4* expression in the oral gastrodermis and in the aboral epidermis of the planula; faint expression is discernable in the tentacle buds, (C) *bmp5-8* is also expressed in the gastrodermis, tentacle buds and aboral epidermis, but appears stronger than *bmp2/4*. (D-D’) *rgm* is strongly expressed in the tentacle buds, siphonoglyph and aboral epidermis. (E-E’) *cv2* is strongly and (F-F’) *admp* is weakly expressed in a cross-like pattern between the tentacles and the in eight epidermal stripes – a pattern identical to the epidermal pSMAD1/5 activity at this stage (see A-A’ and Knabl et al., 2024). tb – tentacle bud, si – siphonoglyph. Scale bars 50 µm (white) and 100 µm (black).

**Figure 2 – Figure Supplement 2.**
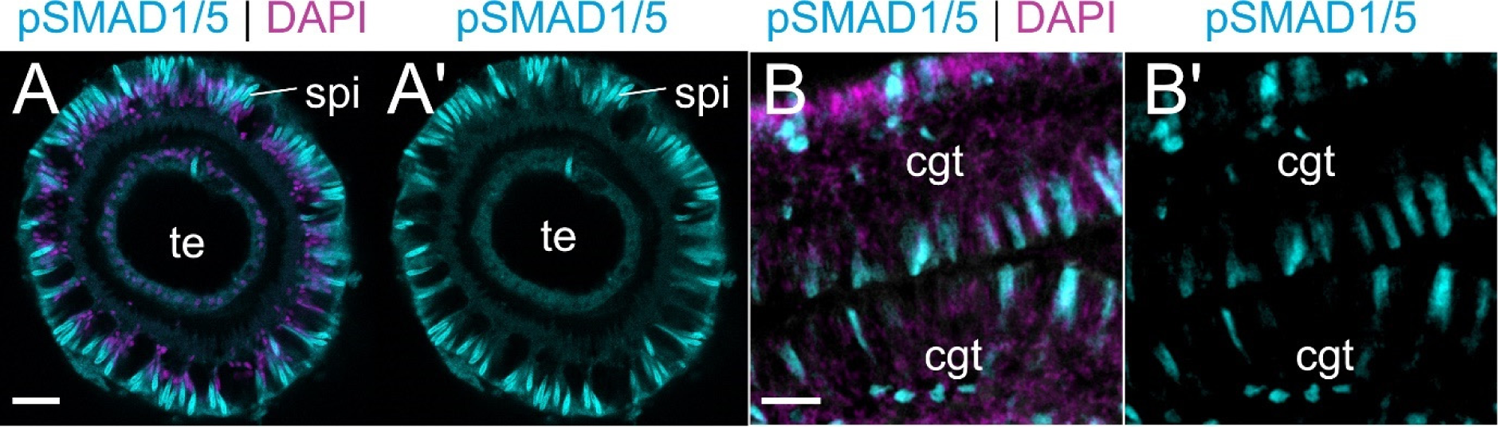
Unspecific staining of spirocytes and muco-glandular cells by pSMAD1/5 antibody. (A-A’) cross-section of a tentacle (te) shows unspecific staining of the coiled thread and capsule wall in the spirocytes (B-B’) close-up of the cnidoglandular tract (cgt) shows unspecific, cytoplasmic signal in muco-glandular cell types. Scale bar 25 µm.

**Figure 2 – Figure Supplement 3.**
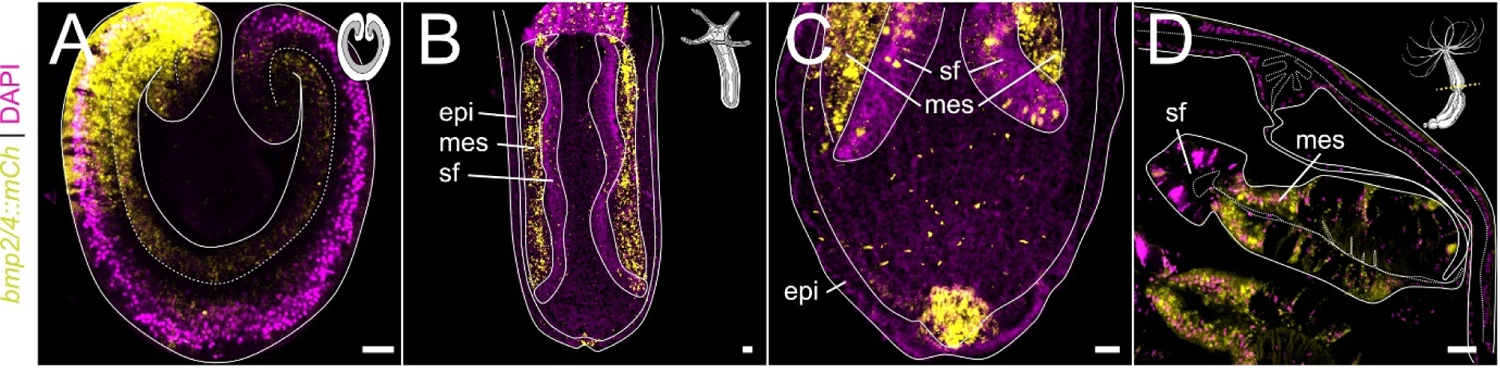
Dynamics of *mCherry* expression in the *bmp2/4::mCh* transgenic reporter line. *bmp2/4::mCherry* expression (yellow) is bilaterally symmetric in the 2d planula larva (A). At primary polyp stage, it is confined to the mesentery gastrodermis (B). Additional aboral expression domain forms in late planula (C, also detectable on B; See also Figure 2 **– Supplement 1** for in situ hybrdization). In adult, expression is observed in the mesentery gastrodermis. epi – epidermis, mes – mesentery, sf - septal filament. Scale bars 50 µm.

**Figure 5 – Figure Supplement 1.**
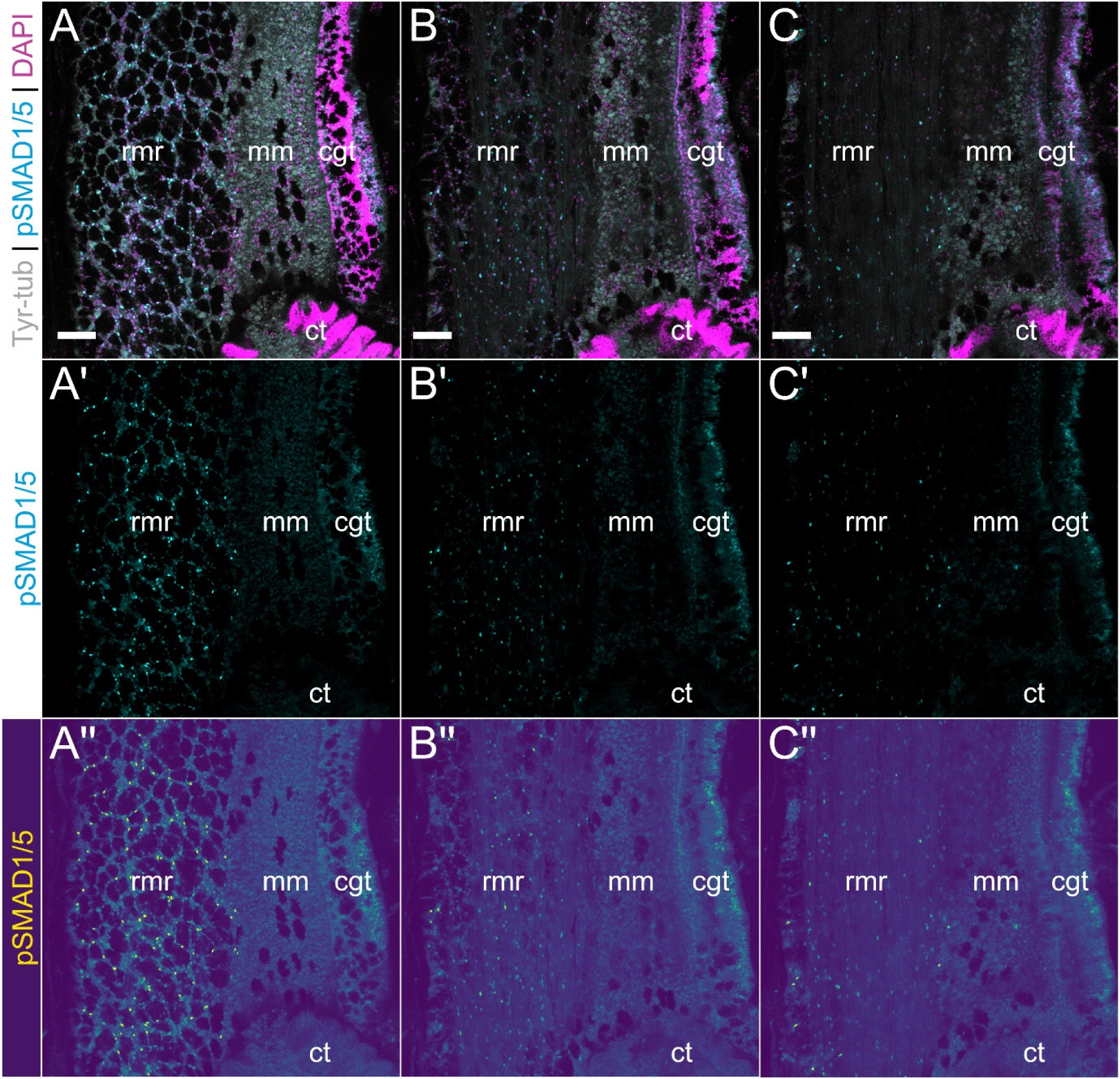
Overview of BMP signaling domains in the mesentery across optical sections. (A-C’’) show different optical sections of the same mesentery region in a longitudinal side view. A’’-C’’ shows the same images as A’-C’ but using a color gradient LUT making the differences in the pSMAD1/5 staining intensity more obvious. Scale bars 50 µm. rmr - retractor muscle region, mm - medial mesentery, cgt - cnidoglandular tract, ct – ciliated tract.

**Figure 6 - Figure Supplement 1.**
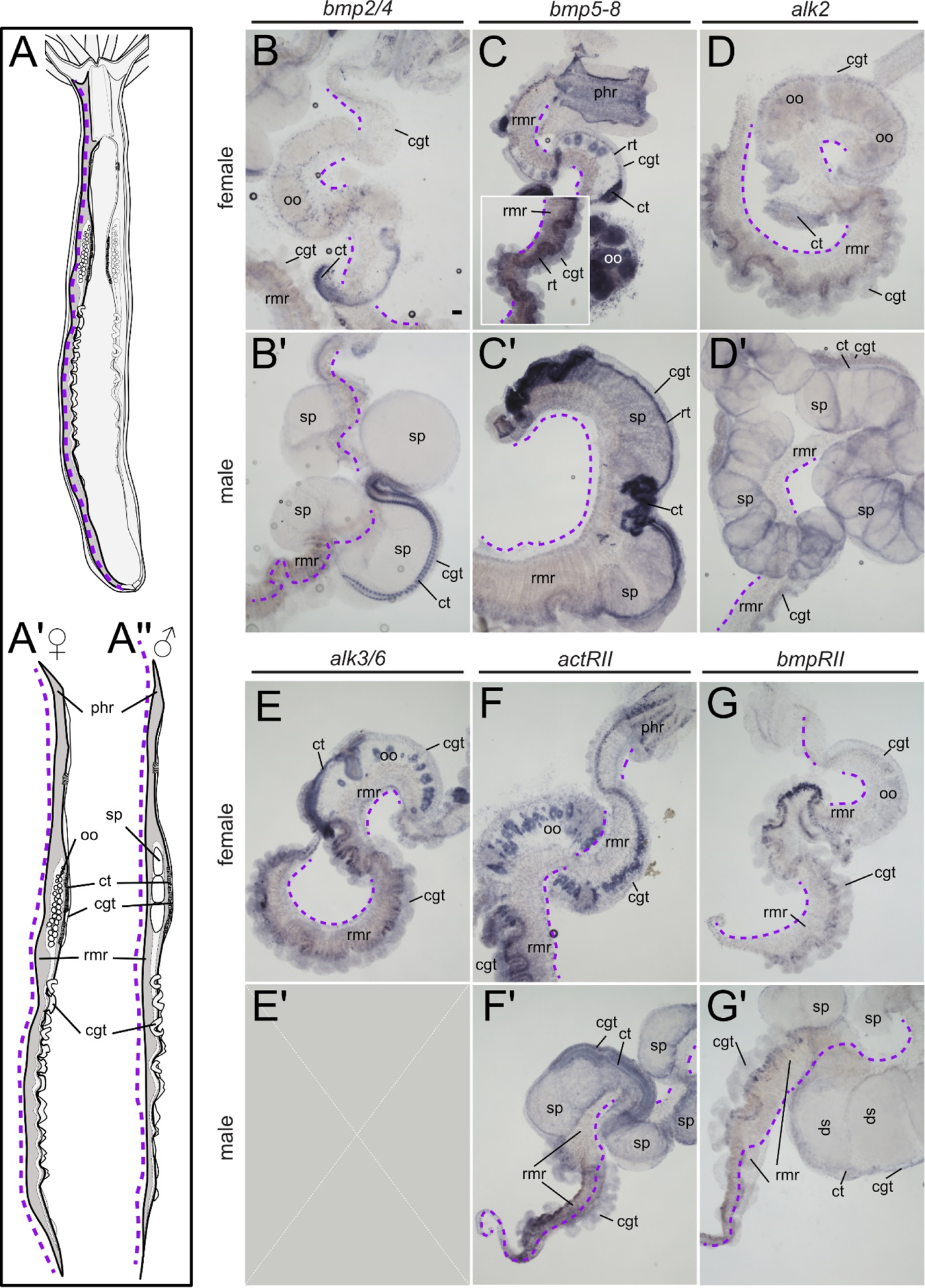
Expression of BMP ligands and receptors in the mesentery. (A) Schematic of a longitudinal section of an adult *Nematostella* polyp and (A’) female and (A’’) male mesentery whole-mounts after dissection, dashed line (purple) indicates the dissection site located between the mesentery stalk (not visible) and the retractor muscle region (rmr). (B-G’) Whole mount tissue pieces of individual female and male mesenteries stained by ISH for the expression of (B-B’) *bmp2/4,* (C-C’) *bmp5-8,* (D-D’) *alk2,* (E-E’) *alk6,* (F-F’) *acrRII* and (G-G’) *bmpRII*. Scale bar 100 µm. phr – pharynx region, rmr – retractor muscle region, cgt – cnidoglandular tract, ct - ciliated tract, oo - oocytes, sp – spermaries,

**Figure 7 – Figure Supplement 1.**
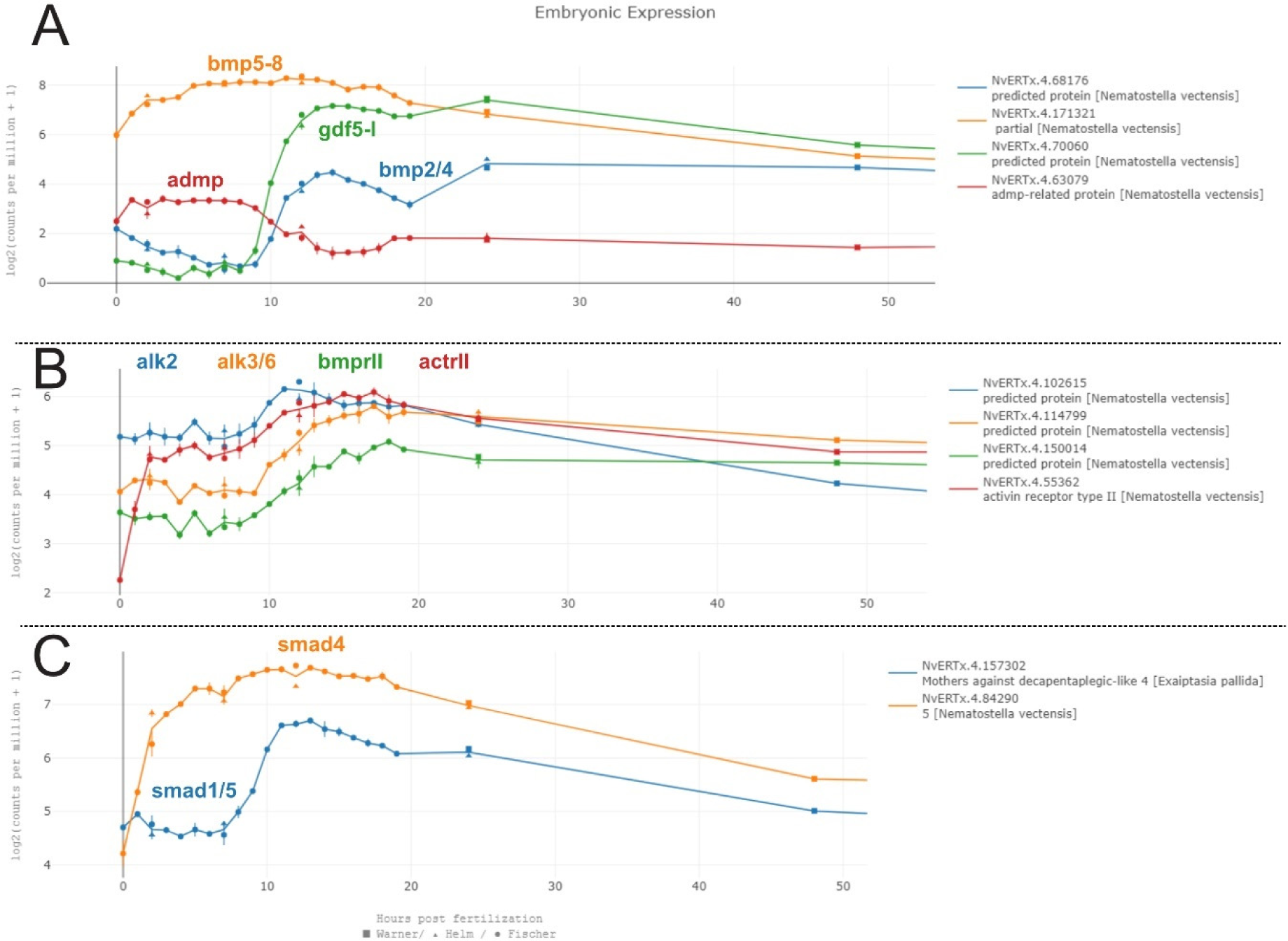
Expression dynamics of BMP pathway genes during embryonic development of *Nematostella*. Expression of (A) *bmp2/4, bmp5-8, gdf5-l, admp,* (B) *alk2, alk6, actRII* and *bmpRII,* (C) *smad1/5* and *smad4* in the egg and during early stages of *Nematostella* development according to the NvERTx database (Fischer & Smith, 2012; Helm et al., 2013; Warner et al., 2018).

**Figure 9 – Figure Supplement 1.**
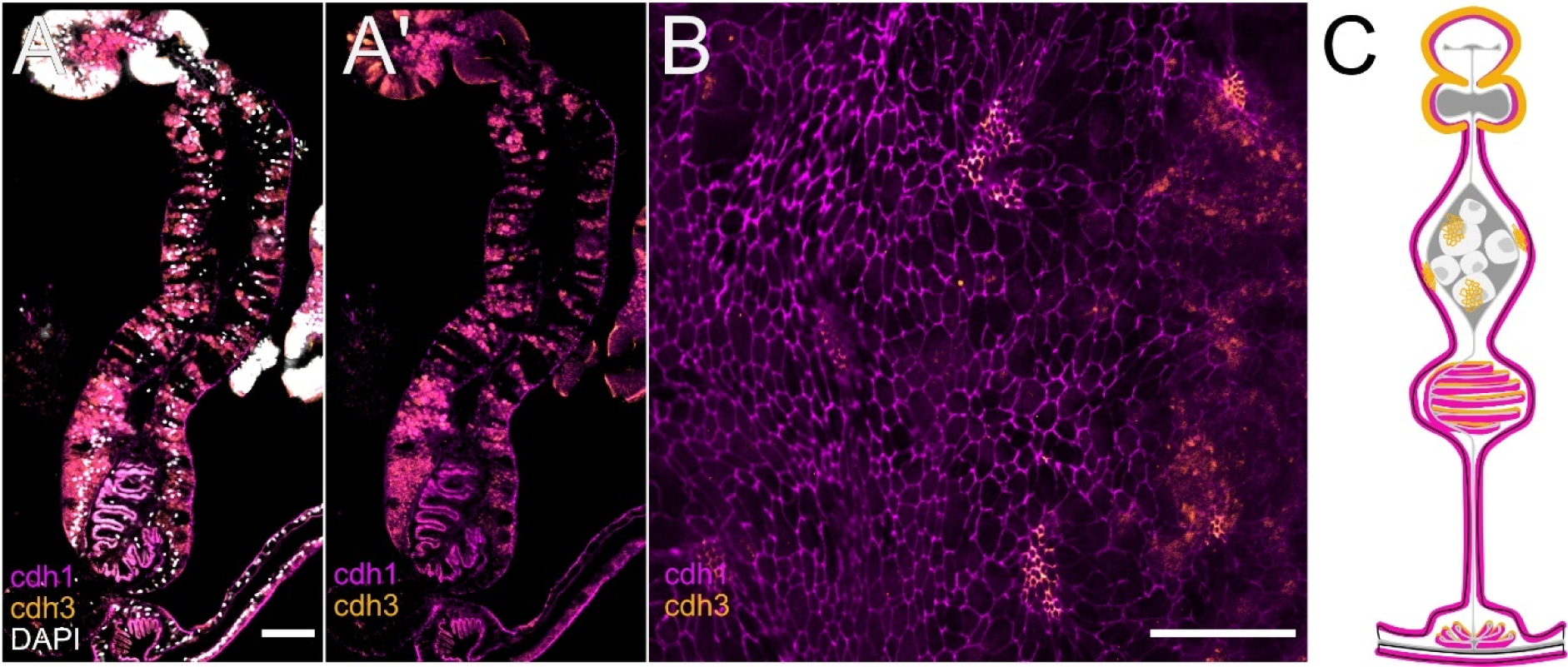
Differential cadherin localization in the mesentery. (A-A’) Broad localization of cdh3 throughout the gastrodermis and distinct localization of cdh1 localization in the septal filament (ingrowth of the pharyngeal tissue (Steinmetz et al., 2017)) and (B) in the accessory cells. Scale bars 50 µm.

**Figure 10 – Figure Supplement 1.**
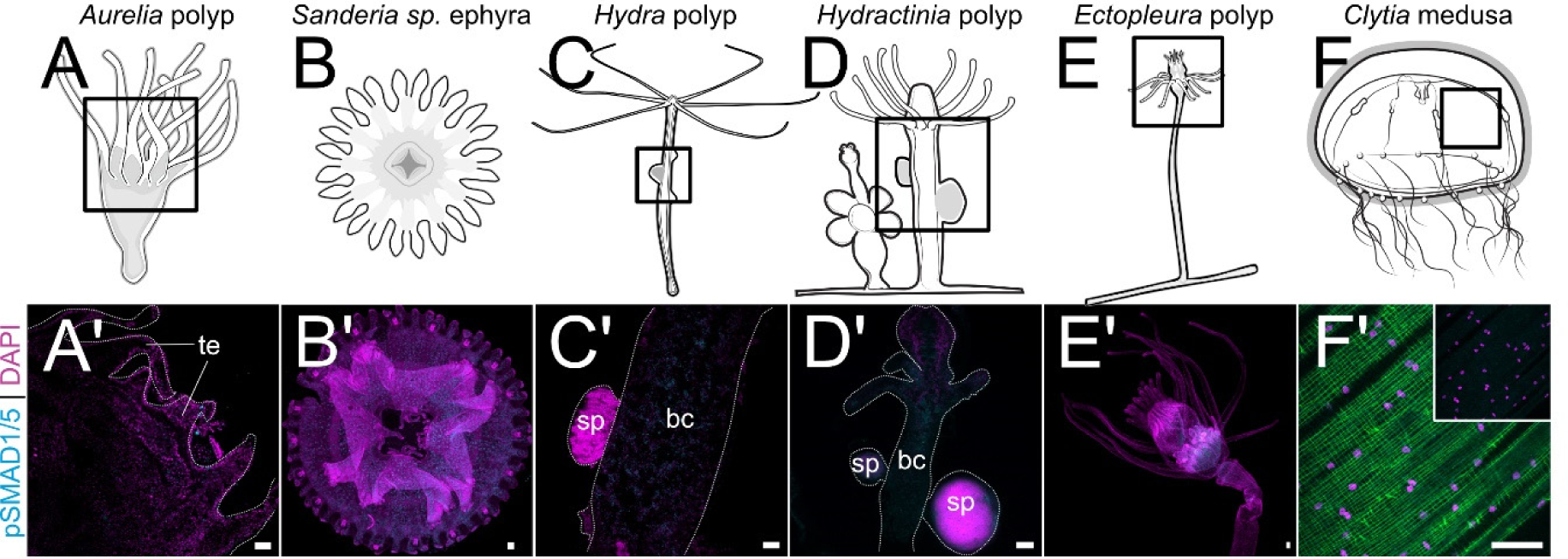
No detection of pSMAD1/5 in several medusozoan cnidarian species. (A’-F’) show no or non-nuclear signal. Scale bar 50 µm.

**Figure 11 – Figure Supplement 1.**
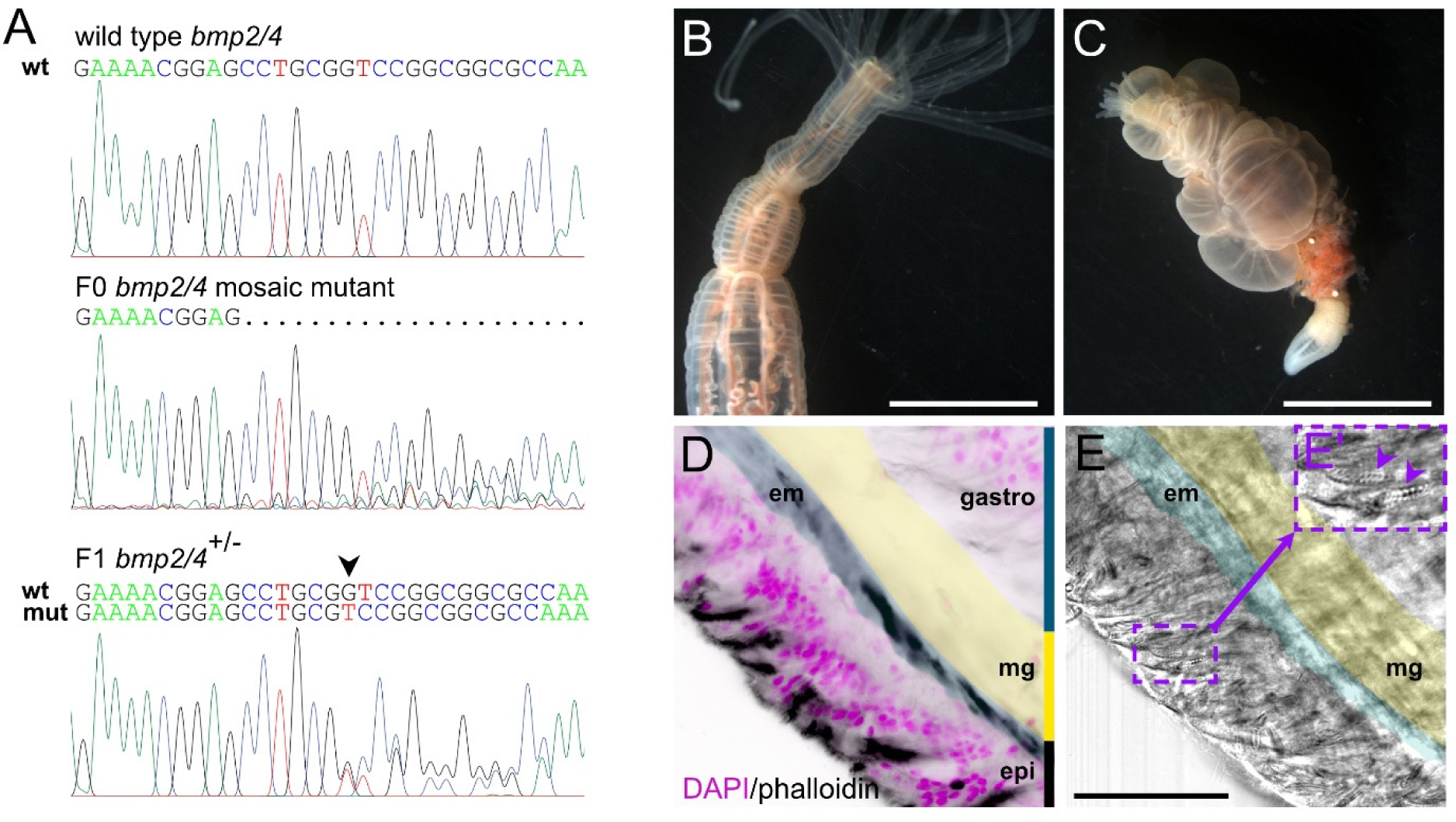
*bmp2/4* knockout animals sporadically develop tentacle-like bubbles. (A) Results of the genotyping of a mosaic F0 mutant and of a F1 heterozygous mutant animal with a single G deletion generating a frame shift. (B-C) Heterozygous *bmp2/4* mutants with a normal morphology (B) and with multiple bubbles (C). (D-E’) Morphological features of the bubble. (D) Phalloidin staining of the bubble tissue shows epidermal muscle fibers exclusive for the normal tentacle tissue. (E-E’) Differential interference contrast image of the same area as on (D) shows multiple spirocysts (arowheads) – a cnidocyte type unique for the tentacles. Scale bars 5 mm (B-C) and 50 µm (E). epi – epidermis, gastro – gastrodermis, mg – mesoglea (yellow shading), em – epidermal muscle (blue shading).

## Footnotes

1 In cnidarian developmental biology literature, the terms „ectoderm“, „endoderm“, and, since the publication of the paper of Steinmetz et al., 2017, also „mesoderm“ are used to describe the germ layers. Since this paper primarily deals with adult tissues, we will be using the morphological terms „epidermis“ for the outer cell layer and „gastrodermis“ for the lining of the gut. We will also extrapolate the use of these terms to embryonic and larval tissues.

## Notes

### Competing Interest Statement

The authors have declared no competing interest.

## References

Bier, E., & De Robertis, E. M. (2015). EMBRYO DEVELOPMENT. BMP gradients: A paradigm for morphogen-mediated developmental patterning. Science, 348(6242), aaa5838. 10.1126/science.aaa5838

Bosch, T. C., & David, C. N. (1987). Stem cells of Hydra magnipapillata can differentiate into somatic cells and germ line cells. Developmental biology, 121(1), 182–191.

Bossert, P. E., Dunn, M. P., & Thomsen, G. H. (2013). A staging system for the regeneration of a polyp from the aboral physa of the anthozoan Cnidarian Nematostella vectensis. Dev Dyn, 242(11), 1320–1331. 10.1002/dvdy.24021

Chen, C. Y., McKinney, S. A., Ellington, L. R., & Gibson, M. C. (2020). Hedgehog signaling is required for endomesodermal patterning and germ cell development in the sea anemone Nematostella vectensis. Elife, 9. 10.7554/eLife.54573

Cole, A. G., Jahnel, S. M., Kaul, S., Steger, J., Hagauer, J., Denner, A., Murguia, P. F., Taudes, E., Zimmermann, B., Reischl, R., Steinmetz, P. R. H., & Technau, U. (2023). Muscle cell-type diversification is driven by bHLH transcription factor expansion and extensive effector gene duplications. Nat Commun, 14(1), 1747. 10.1038/s41467-023-37220-6

Cole, A. G., Steger, J., Hagauer, J., Denner, A., Ferrer Murguia, P., Knabl, P., Narayanaswamy, S., Wick, B., Montenegro, J. D., & Technau, U. (2024). Updated single cell reference atlas for the starlet anemone Nematostella vectensis. Front Zool, 21(1), 8. 10.1186/s12983-024-00529-z

Curantz, C., Doody, C., Horkan, H. R., Krasovec, G., Weavers, P. K., DuBuc, T. Q., & Frank, U. (2024). A positive feedback loop between germ cells and gonads induces and maintains cnidarian sexual reproduction. bioRxiv, 2024.2005.2016.594501. 10.1101/2024.05.16.594501

Denner, A., Steger, J., Ries, A., Morozova-Link, E., Ritter, J., Haas, F., Cole, A. G., & Technau, U. (2023). *Nanos2+* cells give rise to germline and somatic lineages in the sea anemone *Nematostella vectensis*. bioRxiv, 2023.2012.2007.570436. 10.1101/2023.12.07.570436

Driver, I., & Ohlstein, B. (2014). Specification of regional intestinal stem cell identity during Drosophila metamorphosis. Development, 141(9), 1848–1856. 10.1242/dev.104018

DuBuc, T. Q., Schnitzler, C. E., Chrysostomou, E., McMahon, E. T., Febrimarsa, Gahan, J. M., Buggie, T., Gornik, S. G., Hanley, S., Barreira, S. N., Gonzalez, P., Baxevanis, A. D., & Frank, U. (2020). Transcription factor AP2 controls cnidarian germ cell induction. Science, 367(6479), 757–762. 10.1126/science.aay6782

DuBuc, T. Q., Traylor-Knowles, N., & Martindale, M. Q. (2014). Initiating a regenerative response; cellular and molecular features of wound healing in the cnidarian Nematostella vectensis. BMC Biol, 12, 24. 10.1186/1741-7007-12-24

Eckelbarger, K. J., Hand, C., & Uhlinger, K. R. (2008). Ultrastructural features of the trophonema and oogenesis in the starlet sea anemone, Nematostella vectensis (Edwardsiidae). Invertebrate Biology, 127(4), 381–395. 10.1111/j.1744-7410.2008.00146.x

Eckelbarger, K. J., & Larson, R. (1992). Ultrastructure of the ovary and oogenesis in the jellyfish Linuche unguiculata and Stomolophus meleagris, with a review of ovarian structure in the Scyphozoa. Marine Biology, 114(4), 633–643. 10.1007/BF00357260

Extavour, C. G., Pang, K., Matus, D. Q., & Martindale, M. Q. (2005). vasa and nanos expression patterns in a sea anemone and the evolution of bilaterian germ cell specification mechanisms. Evol Dev, 7(3), 201–215. 10.1111/j.1525-142X.2005.05023.x

Finnerty, J. R., Pang, K., Burton, P., Paulson, D., & Martindale, M. Q. (2004). Origins of bilateral symmetry: Hox and dpp expression in a sea anemone. Science, 304(5675), 1335–1337. 10.1126/science.1091946

Fischer, A. H. L., & Smith, J. (2012). Nematostella High-density RNAseq time-course. 10.1575/1912/5981

Friedrichs, M., Wirsdoerfer, F., Flohe, S. B., Schneider, S., Wuelling, M., & Vortkamp, A. (2011). BMP signaling balances proliferation and differentiation of muscle satellite cell descendants. BMC Cell Biol, 12, 26. 10.1186/1471-2121-12-26

Fritzenwanker, J. H., & Technau, U. (2002). Induction of gametogenesis in the basal cnidarian Nematostella vectensis(Anthozoa). Dev Genes Evol, 212(2), 99–103. 10.1007/s00427-002-0214-7

Genikhovich, G., Fried, P., Prunster, M. M., Schinko, J. B., Gilles, A. F., Fredman, D., Meier, K., Iber, D., & Technau, U. (2015). Axis Patterning by BMPs: Cnidarian Network Reveals Evolutionary Constraints. Cell Rep, 10(10), 1646–1654. 10.1016/j.celrep.2015.02.035

Genikhovich, G., & Technau, U. (2009). Induction of spawning in the starlet sea anemone Nematostella vectensis, in vitro fertilization of gametes, and dejellying of zygotes. Cold Spring Harb Protoc, 2009(9), pdb prot5281. 10.1101/pdb.prot5281

Genikhovich, G., & Technau, U. (2017). On the evolution of bilaterality. Development, 144(19), 3392–3404. 10.1242/dev.141507

Gold, D. A., & Jacobs, D. K. (2013). Stem cell dynamics in Cnidaria: are there unifying principles? Dev Genes Evol, 223(1-2), 53–66. 10.1007/s00427-012-0429-1

Gold, D. A., Katsuki, T., Li, Y., Yan, X., Regulski, M., Ibberson, D., Holstein, T., Steele, R. E., Jacobs, D. K., & Greenspan, R. J. (2019). The genome of the jellyfish Aurelia and the evolution of animal complexity. Nat Ecol Evol, 3(1), 96–104. 10.1038/s41559-018-0719-8

He, S., Del Viso, F., Chen, C. Y., Ikmi, A., Kroesen, A. E., & Gibson, M. C. (2018). An axial Hox code controls tissue segmentation and body patterning in Nematostella vectensis. Science, 361(6409), 1377–1380. 10.1126/science.aar8384

Hegarty, S. V., O’Keeffe, G. W., & Sullivan, A. M. (2013). BMP-Smad 1/5/8 signalling in the development of the nervous system. Prog Neurobiol, 109, 28–41. 10.1016/j.pneurobio.2013.07.002

Helm, R. R., Siebert, S., Tulin, S., Smith, J., & Dunn, C. W. (2013). Characterization of differential transcript abundance through time during Nematostella vectensis development. BMC Genomics, 14, 266. 10.1186/1471-2164-14-266

Herrera, A. M., Shuster, S. G., Perriton, C. L., & Cohn, M. J. (2013). Developmental basis of phallus reduction during bird evolution. Curr Biol, 23(12), 1065–1074. 10.1016/j.cub.2013.04.062

Hobmayer, B., Rentzsch, F., & Holstein, T. W. (2001). Identification and expression of HySmad1, a member of the R-Smad family of TGFbeta signal transducers, in the diploblastic metazoan Hydra. Dev Genes Evol, 211(12), 597–602. 10.1007/s00427-001-0198-8

Holstein, T. W. (2023). The Hydra stem cell system - Revisited. Cells Dev, 174, 203846. 10.1016/j.cdev.2023.203846

Houliston, E., Momose, T., & Manuel, M. (2010). Clytia hemisphaerica: a jellyfish cousin joins the laboratory. Trends Genet, 26(4), 159–167. 10.1016/j.tig.2010.01.008

Ikmi, A., Steenbergen, P. J., Anzo, M., McMullen, M. R., Stokkermans, A., Ellington, L. R., & Gibson, M. C. (2020). Feeding-dependent tentacle development in the sea anemone Nematostella vectensis. Nat Commun, 11(1), 4399. 10.1038/s41467-020-18133-0

Itman, C., & Loveland, K. L. (2008). SMAD expression in the testis: an insight into BMP regulation of spermatogenesis. Dev Dyn, 237(1), 97–111. 10.1002/dvdy.21401

Kaltcheva, M. M., Anderson, M. J., Harfe, B. D., & Lewandoski, M. (2016). BMPs are direct triggers of interdigital programmed cell death. Dev Biol, 411(2), 266–276. 10.1016/j.ydbio.2015.12.016

Kayal, E., Bentlage, B., Sabrina Pankey, M., Ohdera, A. H., Medina, M., Plachetzki, D. C., Collins, A. G., & Ryan, J. F. (2018). Phylogenomics provides a robust topology of the major cnidarian lineages and insights on the origins of key organismal traits. BMC Evolutionary Biology, 18(1), 68. 10.1186/s12862-018-1142-0

Khalturin, K., Shinzato, C., Khalturina, M., Hamada, M., Fujie, M., Koyanagi, R., Kanda, M., Goto, H., Anton-Erxleben, F., Toyokawa, M., Toshino, S., & Satoh, N. (2019). Medusozoan genomes inform the evolution of the jellyfish body plan. Nat Ecol Evol, 3(5), 811–822. 10.1038/s41559-019-0853-y

Kirillova, A., Genikhovich, G., Pukhlyakova, E., Demilly, A., Kraus, Y., & Technau, U. (2018). Germ-layer commitment and axis formation in sea anemone embryonic cell aggregates. Proc Natl Acad Sci U S A, 115(8), 1813–1818. 10.1073/pnas.1711516115

Knabl, P., Schauer, A., Pomreinke, A. P., Zimmermann, B., Rogers, K. W., Capek, D., Muller, P., & Genikhovich, G. (2024). Analysis of SMAD1/5 target genes in a sea anemone reveals ZSWIM4-6 as a novel BMP signaling modulator. Elife, 13. 10.7554/eLife.80803

Kraus, J. E., Fredman, D., Wang, W., Khalturin, K., & Technau, U. (2015). Adoption of conserved developmental genes in development and origin of the medusa body plan. Evodevo, 6, 23. 10.1186/s13227-015-0017-3

Krishnapati, L. S., Khade, S., Trimbake, D., Patwardhan, R., Nadimpalli, S. K., & Ghaskadbi, S. (2020). Differential expression of BMP inhibitors gremlin and noggin in Hydra suggests distinct roles during budding and patterning of tentacles. Dev Dyn, 249(12), 1470–1485. 10.1002/dvdy.238

Lauretta, D., Wagner, D., & Penchaszadeh, P. E. (2018). First record of a trophonema in black corals (Cnidaria: Antipatharia). Coral Reefs, 37(2), 581–584. 10.1007/s00338-018-1682-1

Lebouvier, M., Miramon-Puertolas, P., & Steinmetz, P. R. H. (2022). Evolutionarily conserved aspects of animal nutrient uptake and transport in sea anemone vitellogenesis. Curr Biol, 32(21), 4620–4630 e4625. 10.1016/j.cub.2022.08.039

Leclere, L., Horin, C., Chevalier, S., Lapebie, P., Dru, P., Peron, S., Jager, M., Condamine, T., Pottin, K., Romano, S., Steger, J., Sinigaglia, C., Barreau, C., Quiroga Artigas, G., Ruggiero, A., Fourrage, C., Kraus, J. E. M., Poulain, J., Aury, J. M., . . . Copley, R. R. (2019). The genome of the jellyfish Clytia hemisphaerica and the evolution of the cnidarian life-cycle. Nat Ecol Evol, 3(5), 801–810. 10.1038/s41559-019-0833-2

Leclere, L., & Rentzsch, F. (2014). RGM regulates BMP-mediated secondary axis formation in the sea anemone Nematostella vectensis. Cell Rep, 9(5), 1921–1930. 10.1016/j.celrep.2014.11.009

Lewis Ames, C., Ryan, J. F., Bely, A. E., Cartwright, P., & Collins, A. G. (2016). A new transcriptome and transcriptome profiling of adult and larval tissue in the box jellyfish Alatina alata: an emerging model for studying venom, vision and sex. BMC Genomics, 17, 650. 10.1186/s12864-016-2944-3

Lochab, A. K., & Extavour, C. G. (2017). Bone Morphogenetic Protein (BMP) signaling in animal reproductive system development and function. Dev Biol, 427(2), 258–269. 10.1016/j.ydbio.2017.03.002

Matus, D. Q., Pang, K., Marlow, H., Dunn, C. W., Thomsen, G. H., & Martindale, M. Q. (2006). Molecular evidence for deep evolutionary roots of bilaterality in animal development. Proc Natl Acad Sci U S A, 103(30), 11195–11200. 10.1073/pnas.0601257103

Matus, D. Q., Thomsen, G. H., & Martindale, M. Q. (2006). Dorso/ventral genes are asymmetrically expressed and involved in germ-layer demarcation during cnidarian gastrulation. Curr Biol, 16(5), 499–505. 10.1016/j.cub.2006.01.052

Miramón-Puértolas, P., & Steinmetz, P. R. H. (2023). An adult stem-like cell population generates germline and neurons in the sea anemone *Nematostella vectensis*. bioRxiv, 2023.2001.2027.525880. 10.1101/2023.01.27.525880

Moiseeva, E., Rabinowitz, C., Paz, G., & Rinkevich, B. (2017). Histological study on maturation, fertilization and the state of gonadal region following spawning in the model sea anemone, Nematostella vectensis. PLoS One, 12(8), e0182677. 10.1371/journal.pone.0182677

Morsdorf, D., Knabl, P., & Genikhovich, G. (2024). Highly conserved and extremely evolvable: BMP signalling in secondary axis patterning of Cnidaria and Bilateria. Dev Genes Evol. 10.1007/s00427-024-00714-4

Mörsdorf, D., Prünster, M. M., & Genikhovich, G. (2024). Chordin-mediated BMP shuttling patterns the secondary body axis in a cnidarian. bioRxiv, 2024.2005.2027.596067. 10.1101/2024.05.27.596067

Muller, W. A., Teo, R., & Frank, U. (2004). Totipotent migratory stem cells in a hydroid. Dev Biol, 275(1), 215–224. 10.1016/j.ydbio.2004.08.006

Nakanishi, N., Renfer, E., Technau, U., & Rentzsch, F. (2012). Nervous systems of the sea anemone Nematostella vectensis are generated by ectoderm and endoderm and shaped by distinct mechanisms. Development, 139(2), 347–357. 10.1242/dev.071902

Nishimiya-Fujisawa, C., Petersen, H., Koubková-Yu, T. C.-T., Noda, C., Shigenobu, S., Bageritz, J., Fujisawa, T., Simakov, O., Kobayashi, S., & Holstein, T. W. (2023). An ancient split of germline and somatic stem cell lineages in Hydra. bioRxiv, 2023.2007.2004.546637. 10.1101/2023.07.04.546637

Ohdera, A., Ames, C. L., Dikow, R. B., Kayal, E., Chiodin, M., Busby, B., La, S., Pirro, S., Collins, A. G., Medina, M., & Ryan, J. F. (2019). Box, stalked, and upside-down? Draft genomes from diverse jellyfish (Cnidaria, Acraspeda) lineages: Alatina alata (Cubozoa), Calvadosia cruxmelitensis (Staurozoa), and Cassiopea xamachana (Scyphozoa). Gigascience, 8(7). 10.1093/gigascience/giz069

Passamaneck, Y. J., & Martindale, M. Q. (2012). Cell proliferation is necessary for the regeneration of oral structures in the anthozoan cnidarian Nematostella vectensis. BMC Dev Biol, 12, 34. 10.1186/1471-213X-12-34

Praher, D., Zimmermann, B., Genikhovich, G., Columbus-Shenkar, Y., Modepalli, V., Aharoni, R., Moran, Y., & Technau, U. (2017). Characterization of the piRNA pathway during development of the sea anemone Nematostella vectensis. RNA Biol, 14(12), 1727–1741. 10.1080/15476286.2017.1349048

Pukhlyakova, E. A., Kirillova, A. O., Kraus, Y. A., Zimmermann, B., & Technau, U. (2019). A cadherin switch marks germ layer formation in the diploblastic sea anemone Nematostella vectensis. Development, 146(20). 10.1242/dev.174623

Reber-Muller, S., Streitwolf-Engel, R., Yanze, N., Schmid, V., Stierwald, M., Erb, M., & Seipel, K. (2006). BMP2/4 and BMP5-8 in jellyfish development and transdifferentiation. Int J Dev Biol, 50(4), 377–384. 10.1387/ijdb.052085sr

Regan, S. L. P., Knight, P. G., Yovich, J. L., Leung, Y., Arfuso, F., & Dharmarajan, A. (2018). Involvement of Bone Morphogenetic Proteins (BMP) in the Regulation of Ovarian Function. Vitam Horm, 107, 227–261. 10.1016/bs.vh.2018.01.015

Reinhardt, B., Broun, M., Blitz, I. L., & Bode, H. R. (2004). HyBMP5-8b, a BMP5-8 orthologue, acts during axial patterning and tentacle formation in hydra. Dev Biol, 267(1), 43–59. 10.1016/j.ydbio.2003.10.031

Renfer, E., Amon-Hassenzahl, A., Steinmetz, P. R., & Technau, U. (2010). A muscle-specific transgenic reporter line of the sea anemone, Nematostella vectensis. Proc Natl Acad Sci U S A, 107(1), 104–108. 10.1073/pnas.0909148107

Rentzsch, F., Anton, R., Saina, M., Hammerschmidt, M., Holstein, T. W., & Technau, U. (2006). Asymmetric expression of the BMP antagonists chordin and gremlin in the sea anemone Nematostella vectensis: implications for the evolution of axial patterning. Dev Biol, 296(2), 375–387. 10.1016/j.ydbio.2006.06.003

Rentzsch, F., Guder, C., Vocke, D., Hobmayer, B., & Holstein, T. W. (2007). An ancient chordin-like gene in organizer formation of Hydra. Proc Natl Acad Sci U S A, 104(9), 3249–3254. 10.1073/pnas.0604501104

Rentzsch, F., Layden, M., & Manuel, M. (2017). The cellular and molecular basis of cnidarian neurogenesis. Wiley Interdiscip Rev Dev Biol, 6(1). 10.1002/wdev.257

Richards, G. S., & Rentzsch, F. (2014). Transgenic analysis of a SoxB gene reveals neural progenitor cells in the cnidarian Nematostella vectensis. Development, 141(24), 4681–4689. 10.1242/dev.112029

Saina, M., Genikhovich, G., Renfer, E., & Technau, U. (2009). BMPs and chordin regulate patterning of the directive axis in a sea anemone. Proc Natl Acad Sci U S A, 106(44), 18592–18597. 10.1073/pnas.0900151106

Sanders, S. M., Shcheglovitova, M., & Cartwright, P. (2014). Differential gene expression between functionally specialized polyps of the colonial hydrozoan Hydractinia symbiolongicarpus (Phylum Cnidaria). BMC Genomics, 15(1), 406. 10.1186/1471-2164-15-406

Schwaiger, M., Schonauer, A., Rendeiro, A. F., Pribitzer, C., Schauer, A., Gilles, A. F., Schinko, J. B., Renfer, E., Fredman, D., & Technau, U. (2014). Evolutionary conservation of the eumetazoan gene regulatory landscape. Genome Res, 24(4), 639–650. 10.1101/gr.162529.113

Scott, A., & Harrison, P. L. (2009). Gametogenic and reproductive cycles of the sea anemone, Entacmaea quadricolor. Marine Biology, 156(8), 1659–1671. 10.1007/s00227-009-1201-6

Seipel, K., Yanze, N., & Schmid, V. (2004). The germ line and somatic stem cell gene Cniwi in the jellyfish Podocoryne carnea. Int J Dev Biol, 48(1), 1–7. 10.1387/ijdb.15005568

Siebert, S., Farrell, J. A., Cazet, J. F., Abeykoon, Y., Primack, A. S., Schnitzler, C. E., & Juliano, C. E. (2019). Stem cell differentiation trajectories in *Hydra* resolved at single-cell resolution. Science, 365(6451), eaav9314. doi:10.1126/science.aav9314

Steger, J., Cole, A. G., Denner, A., Lebedeva, T., Genikhovich, G., Ries, A., Reischl, R., Taudes, E., Lassnig, M., & Technau, U. (2022). Single-cell transcriptomics identifies conserved regulators of neuroglandular lineages. Cell Rep, 40(12), 111370. 10.1016/j.celrep.2022.111370

Steinmetz, P. R. H., Aman, A., Kraus, J. E. M., & Technau, U. (2017). Gut-like ectodermal tissue in a sea anemone challenges germ layer homology. Nat Ecol Evol, 1(10), 1535–1542. 10.1038/s41559-017-0285-5

Stokkermans, A., Chakrabarti, A., Subramanian, K., Wang, L., Yin, S., Moghe, P., Steenbergen, P., Monke, G., Hiiragi, T., Prevedel, R., Mahadevan, L., & Ikmi, A. (2022). Muscular hydraulics drive larva-polyp morphogenesis. Curr Biol, 32(21), 4707–4718 e4708. 10.1016/j.cub.2022.08.065

Stuart, T., Butler, A., Hoffman, P., Hafemeister, C., Papalexi, E., Mauck, W. M., 3rd, Hao, Y., Stoeckius, M., Smibert, P., & Satija, R. (2019). Comprehensive Integration of Single-Cell Data. Cell, 177(7), 1888–1902 e1821. 10.1016/j.cell.2019.05.031

Tan, E. S., Izumi, R., Takeuchi, Y., Isomura, N., & Takemura, A. (2020). Molecular approaches underlying the oogenic cycle of the scleractinian coral, Acropora tenuis. Sci Rep, 10(1), 9914. 10.1038/s41598-020-66020-x

Tripathi, B. K., & Irvine, K. D. (2022). The wing imaginal disc. Genetics, 220(4). 10.1093/genetics/iyac020

Tucker, R. P., Shibata, B., & Blankenship, T. N. (2011). Ultrastructure of the mesoglea of the sea anemone Nematostella vectensis (Edwardsiidae). Invertebrate Biology, 130(1), 11–24. 10.1111/j.1744-7410.2010.00219.x

Urist, M. R. (1965). Bone: formation by autoinduction. Science, 150(3698), 893–899. 10.1126/science.150.3698.893

Varley, Á., Horkan, H. R., McMahon, E. T., Krasovec, G., & Frank, U. (2023). Pluripotent, germ cell competent adult stem cells underlie cnidarian regenerative ability and clonal growth. Curr Biol, 33(10), 1883–1892.e1883. 10.1016/j.cub.2023.03.039

Vieira, W. A., Wells, K. M., Raymond, M. J., De Souza, L., Garcia, E., & McCusker, C. D. (2019). FGF, BMP, and RA signaling are sufficient for the induction of complete limb regeneration from non-regenerating wounds on Ambystoma mexicanum limbs. Dev Biol, 451(2), 146–157. 10.1016/j.ydbio.2019.04.008

Wang, S., & Chen, Y. G. (2018). BMP signaling in homeostasis, transformation and inflammatory response of intestinal epithelium. Sci China Life Sci, 61(7), 800–807. 10.1007/s11427-018-9310-7

Warner, J. F., Guerlais, V., Amiel, A. R., Johnston, H., Nedoncelle, K., & Rottinger, E. (2018). NvERTx: a gene expression database to compare embryogenesis and regeneration in the sea anemone Nematostella vectensis. Development, 145(10). 10.1242/dev.162867

Watanabe, H., Kuhn, A., Fushiki, M., Agata, K., Ozbek, S., Fujisawa, T., & Holstein, T. W. (2014). Sequential actions of beta-catenin and Bmp pattern the oral nerve net in Nematostella vectensis. Nat Commun, 5, 5536. 10.1038/ncomms6536

Watanabe, H., Schmidt, H. A., Kuhn, A., Hoger, S. K., Kocagoz, Y., Laumann-Lipp, N., Ozbek, S., & Holstein, T. W. (2014). Nodal signalling determines biradial asymmetry in Hydra. Nature, 515(7525), 112–115. 10.1038/nature13666

Wedi, S. E., & Dunn, D. F. (1983). Gametogenesis and Reproductive Periodicity of the Subtidal Sea Anemone Urticina Lofotensis (Coelenterata: Actiniaria) in California. Biol Bull, 165(2), 458–472. 10.2307/1541212

Wenger, Y., Buzgariu, W., Perruchoud, C., Loichot, G., & Galliot, B. (2019). Generic and context-dependent gene modulations during *Hydra* whole body regeneration. bioRxiv, 587147. 10.1101/587147

Wu, M., Wu, S., Chen, W., & Li, Y. P. (2024). The roles and regulatory mechanisms of TGF-beta and BMP signaling in bone and cartilage development, homeostasis and disease. Cell Res, 34(2), 101–123. 10.1038/s41422-023-00918-9

Zhang, Y., & Que, J. (2020). BMP Signaling in Development, Stem Cells, and Diseases of the Gastrointestinal Tract. Annu Rev Physiol, 82, 251–273. 10.1146/annurev-physiol-021119-034500

Zou, H., & Niswander, L. (1996). Requirement for BMP signaling in interdigital apoptosis and scale formation. Science, 272(5262), 738–741. 10.1126/science.272.5262.738

